# The carboxyl-terminal sequence of Bim enables Bax activation and killing of unprimed cells

**DOI:** 10.1101/554907

**Authors:** Xiaoke Chi, Dang Nguyen, James M Pemberton, Elizabeth J Osterlund, Qian Liu, Hetal Brahmbhatt, Zhi Zhang, Jialing Lin, Brian Leber, David W. Andrews

**Author notes:** These authors contributed equally to this work (Dang Nguyen, James M Pemberton). **Lead Contact:** Dr. David W Andrews Address: Sunnybrook Research Institute, Room M7-621, 2075 Bayview Avenue, Toronto, ON, Canada, M4N 3M5.

## Abstract

The Bcl-2 family BH3 protein Bim promotes apoptosis at mitochondria by activating the pore forming proteins Bax and Bak and by inhibiting the anti-apoptotic proteins Bcl-XL, Bcl-2 and Mcl-1. Bim binds to these proteins via its BH3 domain and to the mitochondrial membrane by a carboxyl-terminal sequence (CTS). In cells killed by Bim, the expression of a Bim mutant in which the CTS was deleted (BimL-dCTS) triggered variable amounts of apoptosis that correlated with inhibition of anti-apoptotic proteins being sufficient to permeabilize mitochondria isolated from the same cells. Detailed analysis of the molecular mechanism demonstrated that BimL-dCTS inhibited Bcl-XL but did not activate Bax. Our examination of additional point mutants unexpectedly revealed that the CTS of Bim is required for physiological concentrations of Bim to activate Bax and that different residues in the CTS enable Bax activation and binding to membranes.

## Introduction

Apoptosis is a highly conserved form of programmed cell death that can be triggered by extrinsic or intrinsic signals. It plays a fundamental role in maintaining homeostasis by eliminating old, excessive or dysfunctional cells in multi-cellular organisms (Kerr, Wyllie, and Currie 1972). Defective regulation of apoptosis has been found in many diseases (Favaloro et al. 2012) and is considered one of the hallmarks of cancer (Hanahan and Weinberg 2011).

Bcl-2 family proteins play a decisive role in apoptosis initiated by intrinsic signaling by regulating the integrity of the mitochondrial outer membrane (MOM). Commitment to apoptosis is generally regarded as due to MOM permeabilization (MOMP) releasing cytochrome c and pro-apoptotic factors from the intermembrane space into the cytoplasm. These factors activate the executioner caspases that mediate cell death (Chipuk, Bouchier-Hayes, and Green 2006). Direct interactions between Bcl-2 family proteins govern both initiation and inhibition of MOMP (Kale, Osterlund, and Andrews 2017). The Bcl-2 family of proteins that regulate apoptosis includes the anti-apoptotic proteins Bcl-XL, Bcl-2 and Mcl-1 that inhibit the process and share four Bcl-2 homology domains. These homology domains, referred to as BH domains, are also shared by the pro-apoptotic proteins Bax and Bak that permeabilize the MOM directly. Both pro- and anti-apoptotic multi-domain Bcl-2 family proteins are regulated by direct binding interactions with a group of proteins including Bim, Bid, Puma, Hrk, Bad and Noxa that contain a single region of homology, the Bcl-2 homology domain number 3, and are therefore referred to collectively as BH3-proteins. These proteins promote apoptosis by releasing sequestered activated Bax, Bak and BH3-proteins that activate Bax and Bak from one or more of the anti-apoptotic proteins. The subset of BH3 proteins that bind to and activate Bax or Bak include Bid, Bim and Puma(Chi et al. 2014). Thus far, the biochemical basis for the differences between BH3-proteins that inhibit anti-apoptotic proteins and those that activate Bax and Bak has been attributed entirely to differences in affinities of the BH3-domain for the BH3-peptide binding sites on multi-domain pro- and anti-apoptotic proteins. However, static affinities and variations in expression levels permit only coarse regulation of cell death. Changes in the equilibrium binding of Bcl-2 family proteins on the MOM enable finer control. For example, at physiologic concentrations the BH3 protein Bid only activates Bax after Bid has bound to a membrane and undergone a specific conformational change (Lovell et al. 2008; Shamas-Din, Bindner, et al. 2013). Binding to membranes also enables interaction of Bid with MTCH2 on the MOM to greatly accelerate the Bid conformational change that results in Bax activation (Shamas-Din, Bindner, et al. 2013). However, it remains unclear whether membrane interactions by other BH3 proteins like Bim contribute to Bax activation.

The BH3-protein Bim is an important mediator of apoptosis initiated by many intracellular stressors (Concannon et al. 2010; Mahajan et al. 2014; Puthalakath et al. 2007). Three major isoforms of Bim result from alternative mRNA splicing: BimEL, BimL, and BimS (O’Connor et al. 1998). All three isoforms include the BH3-domain required for binding other Bcl-2 family proteins, and a C-terminal sequence (CTS) that binds the protein to the MOM (Wilfling et al. 2012). BimEL and BimL also share a dynein light chain binding motif (LC1) that sequesters these isoforms at the cytoskeleton (Lei and Davis 2003). The absence of this LC1 binding motif in BimS likely accounts for the constitutive activity of the isoform in cells (Lei and Davis 2003). Accordingly, BimS is regulated transcriptionally and rarely present in healthy cells, while BimL and BimEL are present in most tissue types (O’Reilly et al. 2000). Bim has a particularly important function as a regulator of anti-apoptotic proteins, as it binds and thereby inhibits by mutual sequestration all known anti-apoptotic proteins (Chen et al. 2005; Shamas-Din, Kale, et al. 2013). Until recently, It was unknown why Bim binds to Bcl-XL with sufficient affinity to resist displacement by small molecule BH3-mimetics, while other BH3 proteins, such as Bad, are easily displaced (Aranovich et al. 2012). In addition to interactions via the BH3-domain, residues within the Bim CTS bind to Bcl-XL, and thereby increase the affinity of the interaction by “double-bolt locking” providing an explanation for the observations with BH3 mimetic drugs (Liu et al. 2019). Here we investigated whether the CTS of Bim also contributes to the functional and physical interactions between Bim and Bax.

We demonstrate that both primary cells and cell lines have a range of apoptotic responses to the expression of a truncated BimL protein lacking the CTS (BimL-dCTS), while expression of full-length BimL was sufficient to kill all of these cells. To determine the molecular mechanism that underlies this difference, the two pro-apoptotic functions of Bim; activation of Bax and inhibition of Bcl-XL, were quantified using purified full-length BimL protein and cell free assays. Replacing the CTS of Bim with an alternative tail-anchor that binds the protein to mitochondrial membranes did not fully restore Bax activation function, demonstrating that sequences within the Bim CTS rather than membrane binding contribute to Bax activation. Site-directed mutagenesis of the Bim CTS also revealed residues important for binding to membranes that were not required for Bax activation (e.g. I125). Furthermore, specific residues within the CTS were identified that are required for BimL to efficiently activate Bax, but that are not required for BimL to bind to and inhibit Bcl-XL. Evidence in cell free assays demonstrated that BimL CTS residues L129 and I132 physically interact with Bax and are required to activate it. These mutants were used to show that BimL residues L129 and I132 are also required for BimL to efficiently kill cells resistant to BimL-dCTS, demonstrating that it is necessary to activate Bax to kill these unprimed cells. Together, our data demonstrates that the unusual sequence of the CTS of Bim separately controls both membrane binding and Bax activation.

## Results

### The CTS of Bim variably contributes to the pro-apoptotic activity of Bim in different cell lines

Removing the CTS from Bim abrogates pro-apoptotic activity in HEK293 cells (Weber et al. 2007). While this observation has generally been ascribed to loss of binding of Bim to MOM our observation that the CTS is also involved in binding BimEL to Bcl-XL (Liu et al. 2019) suggested that there may be other explanations for the loss of pro-apoptotic activity for Bim when the CTS is removed. To determine the contribution of the Bim CTS to pro-apoptotic activity, a BimL mutant was generated in which the previously characterized membrane binding domain (carboxyl-terminal residues P121-H140) were deleted (BimL-dCTS) (Wilfling et al. 2012), (Liu et al. 2019). This mutant was expressed in cells and the effectiveness of induction of cell death was compared to expression of full-length BimL by confocal microscopy. To detect expression of the constructs in live cells, they included an N-terminally fused Venus fluorescent protein (indicated by a superscripted v in the name). Thus a construct in which Venus was fused to BimL is referred to here as ^V^BimL while the mutant lacking the CTS is ^V^BimL-dCTS. As an inactive control we used ^V^BimL-4E a mutant in which four conserved hydrophobic residues in the BH3 domain of BimL were replaced with glutamate, thereby preventing binding to all other Bcl-2 family proteins (Chen et al. 2005), (Liu et al. 2019).

To assay pro-apoptotic activity, the constructs were expressed in primary cells and cell lines and both expression and cell death were measured using confocal microscopy. Apoptosis was assessed by detecting externalization of phosphatidylserine by Annexin V staining in cells expressing detectable levels of ^V^BimL or the ^V^BimL mutants as measured by Venus fluorescence. As expected, expression of ^V^BimL induced apoptosis in all cell types tested, while the negative control ^V^BimL-4E did not (Figure 1A). As reported previously for Bim-dCTS, the fluorescent version (^V^Bim-dCTS) failed to induce cell death in HEK293 cells (Weber et al. 2007). In contrast, expression of ^V^BimL-dCTS induced apoptosis to levels similar to ^V^BimL in HCT116 and MEF cells but had reduced potency in BMK and CAMA-1 cells. Thus the CTS of Bim contributed variably to the pro-apoptotic activity of Bim in different cell lines despite having equal expression across all cell types (Figure 1 – figure supplement 1).

**Figure 1:**
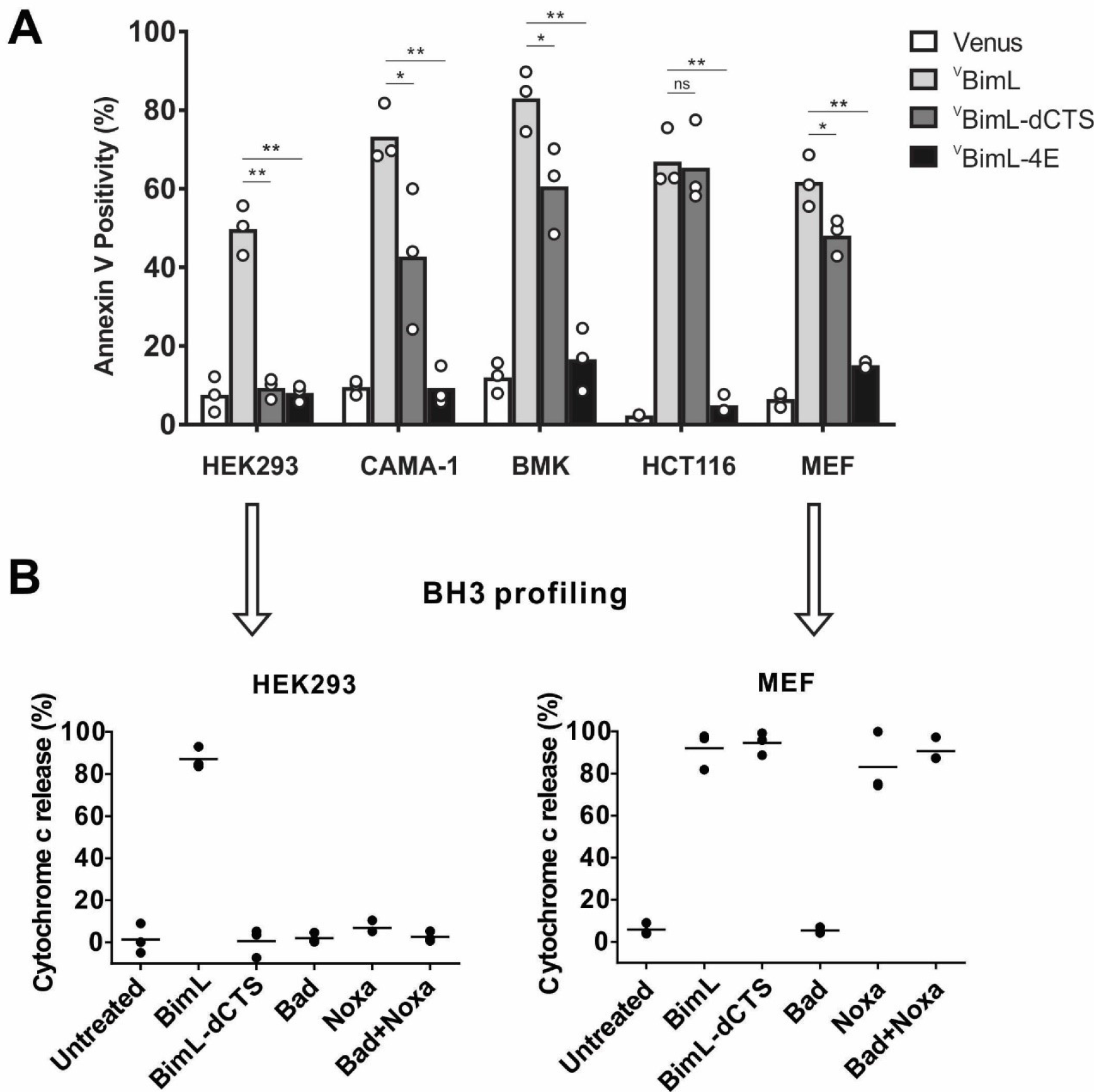
Cell lines demonstrate a range of apoptotic response to BimL-dCTS expression. (A) Venus, ^V^BimL, ^V^BimL-dCTS or ^V^Bim-4E were expressed in the Indicated cell lines by transient transfection. The cells were stained with the nuclear dye Draq5 and rhodamine labelled Annexin V, and imaged by confocal microscopy to identify cells undergoing apoptosis. At least 400 cells were analyzed for each condition. The Y axis indicates the percentage of Venus positive cells that stained positive with Annexin V. Open circles represent the average for each replicate, while the bar height, relative to the y-axis, represents the average for all three replicates. The means were assessed for significant differences using a one-way ANOVA within each group followed by a Tukey’s multiple comparisons test. *p-values less than 0.05, **p-values less than 0.01, ns are non-significant p-values (>0.05). (B) BH3 profiling of mitochondria isolated from HEK293 and MEF cells. Mitochondria (1mg/mL) were incubated with 500 nM of the indicated recombinant BH3 protein(s) for 1 hour at 37°C. Cytochrome c release, indicative of MOMP, was quantified by immunoblotting. Data from three independent experiments are shown as individual points, with lines representing the average. Some dots are not visible due to overlap.

To determine if this difference in response to ^V^BimL-dCTS expression is a function of the extent to which the apoptotic machinery is loaded in mitochondrial outer-membranes, mitochondria were purified from cells resistant (HEK293) and sensitive (MEF) to ^V^BimL-dCTS expression and assayed by BH3-profiling (Potter and Letai 2016) to measure loading of anti-apoptotic proteins with BH3 proteins or active Bax/Bak. Unlike BH3-profiling experiments conducted with BH3-peptides, in these experiments purified full-length proteins were used. Thus, purified BimL, BimL-dCTS, Bad and Noxa proteins were incubated with mitochondria from each of the cell lines and mitochondrial outer membrane permeabilization (MOMP) was measured by separating supernatant and pellet fractions for each reaction, and immunoblotting for cytochrome c released from the intermembrane space as previously described (Pogmore et al. 2016). Immunoblots were quantified and MOMP assessed as % cytochrome c released (Figure 1B). As expected from the data in Figure 1A, addition of recombinant BimL was sufficient to induce cytochrome c release from mitochondria from both HEK293 and MEF cells. However, addition of BimL-dCTS induced cytochrome c release only in the MEF mitochondria confirming that resistance to BimL-dCTS in HEK293 cells is manifest at mitochondria.

One potential explanation for this difference is that the mitochondria in the cell lines have different dependencies on multi-domain anti-apoptotic proteins for survival, a phenomenon known as priming. If BimL-dCTS has lost one of the functions of Bim such as activating Bax or Bak or inhibiting one of the multi-domain anti-apoptotic proteins Bcl-2, Bcl-XL and Mcl-1 it would be expected to have different activities on mitochondria with different priming. Therefore, to better understand why BimL-dCTS can only permeabilize MEF mitochondria and not mitochondria from HEK293 cells, we compared the sensitivity of mitochondria from the two cell types to addition of BH3-proteins Bad and Noxa that inhibit Bcl-2 and Bcl-XL or Mcl-1, respectively, but that do not activate Bax or Bak (Kale, Osterlund, and Andrews 2017). Incubation of full length Bad and/or Noxa with mitochondria from HEK293 cells failed to induce cytochrome c release, while the addition of Noxa was sufficient to permeabilize MEF mitochondria (Figure 1B). This data suggests that HEK293 cells do not depend on expression of Bcl-2, Bcl-XL or Mcl-1 sequestering active Bax, Bak or their BH3-activators while mitochondria from MEFs depend on expression of Mcl-1 to prevent apoptosis (Lessene et al. 2013). The results further suggest that removal of the CTS from BimL results in a mutant protein that only kills cells dependent on one or more multi-domain anti-apoptotic proteins for survival. That BimL-dCTS does not kill HEK293 cells further suggests that it does not activate sufficient Bax or Bak to overcome the unoccupied anti-apoptotic proteins in this cell line. In this way BimL-dCTS functions as a sensitizer similar to proteins like Bad and Noxa. However, unlike other relatively specific sensitizer proteins, the known binding activity of the BH3 region of BimL-dCTS suggests that it inhibits Bcl-2, Bcl-XL and Mcl-1. Indeed we have shown that in live cells BimEL-dCTS binds to Bcl-2 and Bcl-XL but is more easily displaced than BimEL by small molecule BH3 mimetics (Liu et al. 2019).

### Full-length BimL is required to kill cultures of primary cortical neurons

Our data with cell lines and their respective purified mitochondria suggests that BimL-dCTS does not kill cells that do not depend on anti-apoptotic proteins for survival. To test this in a more biologically relevant system, we cultured primary murine cortical neurons and assayed their response to expression of the BimL mutants. For regulated expression in primary cortical neurons the coding regions for ^V^BimL, ^V^BimL-4E, and ^V^BimL-dCTS were cloned into a tetracycline-responsive lentiviral vector, and introduced into primary cortical neuron cultures through lentiviral infection. After culture for 8 days *in vitro*, BimL expression was induced in the neurons by the addition of doxycycline. Neuronal cell death was assayed using confocal microscopy after staining neurons with propidium iodide (PI), a dye that exclusively stains the nuclei of dead cells. Quantification of Venus-expressing neuronal cell bodies revealed that as expected ^V^BimL expression killed cultured primary neurons while ^V^BimL-4E did not (Figure 2A-B). However, the expression of ^V^BimL-dCTS was largely ineffective to induce cell death in cultured primary cortical neurons (Figure 2B). Our data is consistent with previous reports suggesting that primary murine cultures of hippocampal neurons become resistant to induction of apoptosis by external stimuli over time in culture. This resistance has been reported to be due to a difference in Bcl-2 family protein expression that results in decreased mitochondrial ‘priming’, explaining why our cultures of primary cortical neurons are resistant to ^V^BimL-dCTS (Sarosiek et al. 2016).

**Figure 2:**
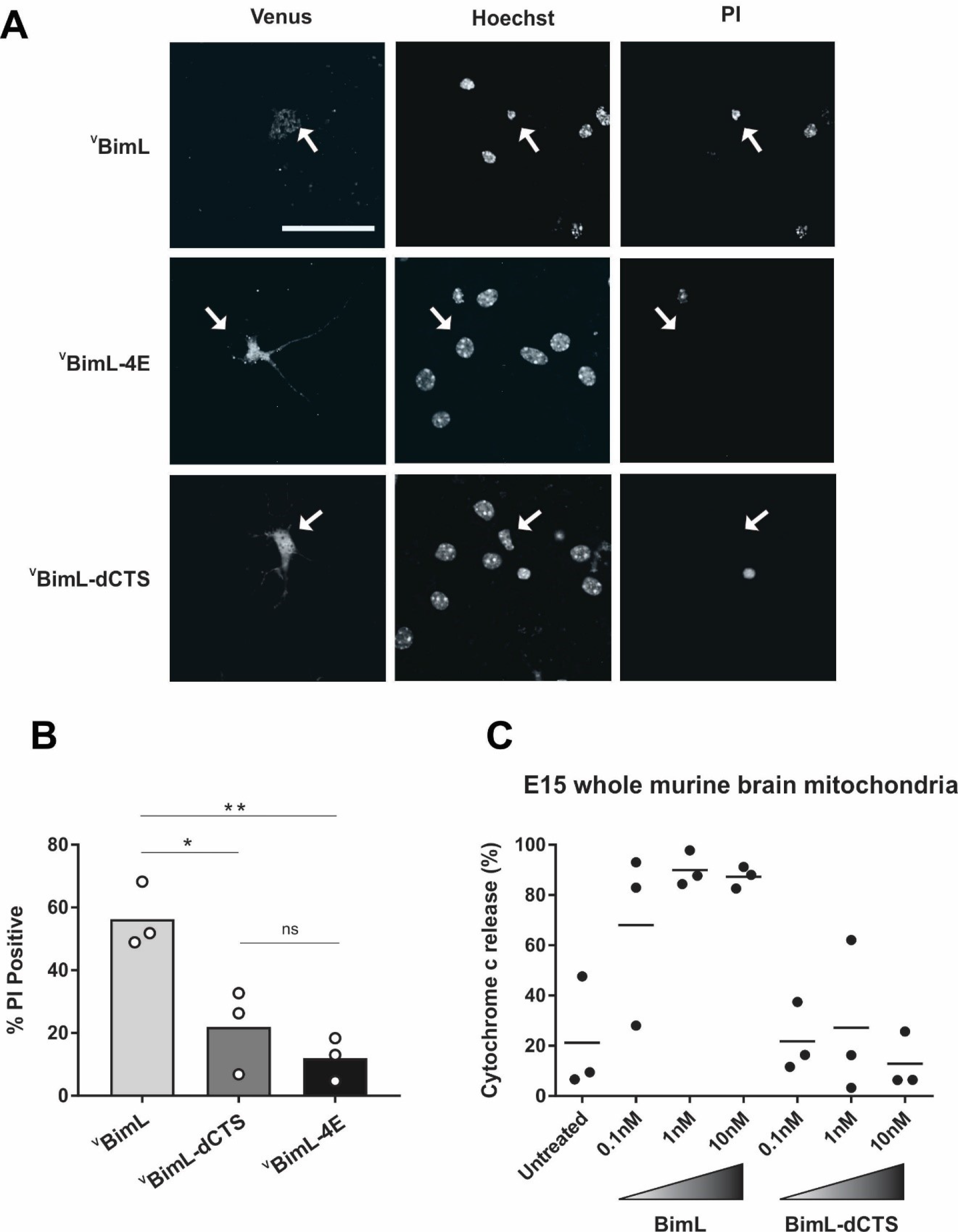
Full-length BimL is required to kill cultures of primary cortical neurons. (A) Representative images of primary cortical neurons infected with lentivirus to express ^V^BimL, ^V^BimL-dCTS or ^V^BimBH3-4E. White arrows indicate neurons expressing Venus fluorescence. Scale bar is 80μm. B) Quantified data from Venus expressing primary cortical neurons. Percentage of Venus expressing cells with nuclei PI intensity scores above threshold (% PI Positive). Open circles; averages for three biological replicates each representing 90-1000 cells analyzed. The Bar height; mean. A one-way ANOVA was used followed by a Tukey’s multiple comparisons test to compare the means of each group. * p< 0.05; ** p,0.01; ns p>0.05. (C) Mitochondria extracted from embryonic day 15 (E15) mouse brains (0.5 mg/mL) were incubated with the indicated BH3-only proteins. Cytochrome c release, indicative of MOMP, was quantified using immunoblotting. Each point (black circle) represents one independent replicate, with the line representing the average across all three.

To determine if resistance to induction of cell death by BimL-dCTS is due to differential sensitivity of neuronal mitochondria to induction of MOMP by BimL and BimL-dCTS, mitochondria were isolated from embryonic day 15 (E15) mouse brains, the same age used to culture primary cortical neurons. Brain mitochondria were used instead of isolating mitochondria from neuronal cultures due to the low yield from primary cultured neurons. Untreated mitochondria from day E15 brain released only low levels of cytochrome c. As expected, addition of 0.1nM recombinant BimL was sufficient to elicit MOMP as measured by cytochrome c release and detection in the supernatant. In contrast, 100 times more BimL-dCTS (10nM) failed to induce MOMP (Figure 2C).

Taken together our data suggest that BimL-dCTS kills cells in which the mitochondria are sensitive to inhibition of anti-apoptotic proteins by sensitizers such as Bad and Noxa. Thus, BimL-dCTS did not permeabilize mitochondria extracted from HEK293 cells or E15 whole murine brains, and as a result, BimL-dCTS expression did not kill HEK293 cells or primary cultures of cortical neurons. This finding suggests that inhibition of anti-apoptotic proteins is not sufficient to kill these cells. Therefore, BimL-dCTS differs mechanistically from BimL as the latter kills both cell types resistant and sensitive to BimL-dCTS. Compared to BimL, BimL-dCTS is missing the membrane binding domain and therefore is not expected to localize at mitochondria (Liu et al. 2019), however, the relationship between Bim binding to membranes and Bim mediated Bax activation has not been extensively studied. To determine how the molecular mechanism of BimL-dCTS differs from BimL the activities of the proteins were analyzed using cell free assays.

### The Bim CTS mediates BimL binding to both Bax and membranes

To investigate the pro-apoptotic mechanism of BimL and BimL-dCTS without interference from other cellular components, both were purified as full-length recombinant proteins and assayed using liposomes and/or isolated mitochondria. To measure direct-activation of Bax by Bim, either BimL or BimL-dCTS was incubated with recombinant Bax and liposomes encapsulating the dye and quencher pair: ANTS (8-Aminonaphthalene-1,3,6-Trisulfonic Acid, Disodium Salt) and DPX (*p*-Xylene-Bis-Pyridinium Bromide). In this well-established assay (Kale et al. 2014), increasing amounts of BimL activated Bax resulting in membrane permeabilization measured as an increase in fluorescence due to the release and separation of encapsulated dye and quencher (Figure 3A). This result is consistent with previous observations that picomolar concentrations of BimL induce Bax-mediated membrane permeabilization (Sarosiek et al. 2013). In contrast, three orders of magnitude higher concentrations of BimL-dCTS were required to induce Bax-mediated liposome permeabilization (Figure 3A), suggesting that either or both of binding to membranes and the specific CTS of Bim are required for efficient Bax activation. As expected, similar results were obtained for Bax-mediated release of mitochondrial intermembrane space proteins (Figure 3B). For these experiments, MOMP was measured as release of the fluorescent protein mCherry fused to the N-terminal mitochondrial import signal of SMAC (SMAC-mCherry) from the intermembrane space of mitochondria (Shamas-Din et al. 2014). Similar to the results with liposomes (Figure 3A), and mitochondria from cell lines (Figures 1-2) BimL but not BimL-dCTS triggered Bax mediated SMAC-mCherry release from mitochondria isolated from Bax −/− Bak−/− cells (Figure 3B). In experiments with liposomes and mitochondria, very small amounts of Bim were sufficient to trigger membrane permeabilization because once activated, Bax recruits and activates additional Bax molecules (Tan et al. 2006).

**Figure 3:**
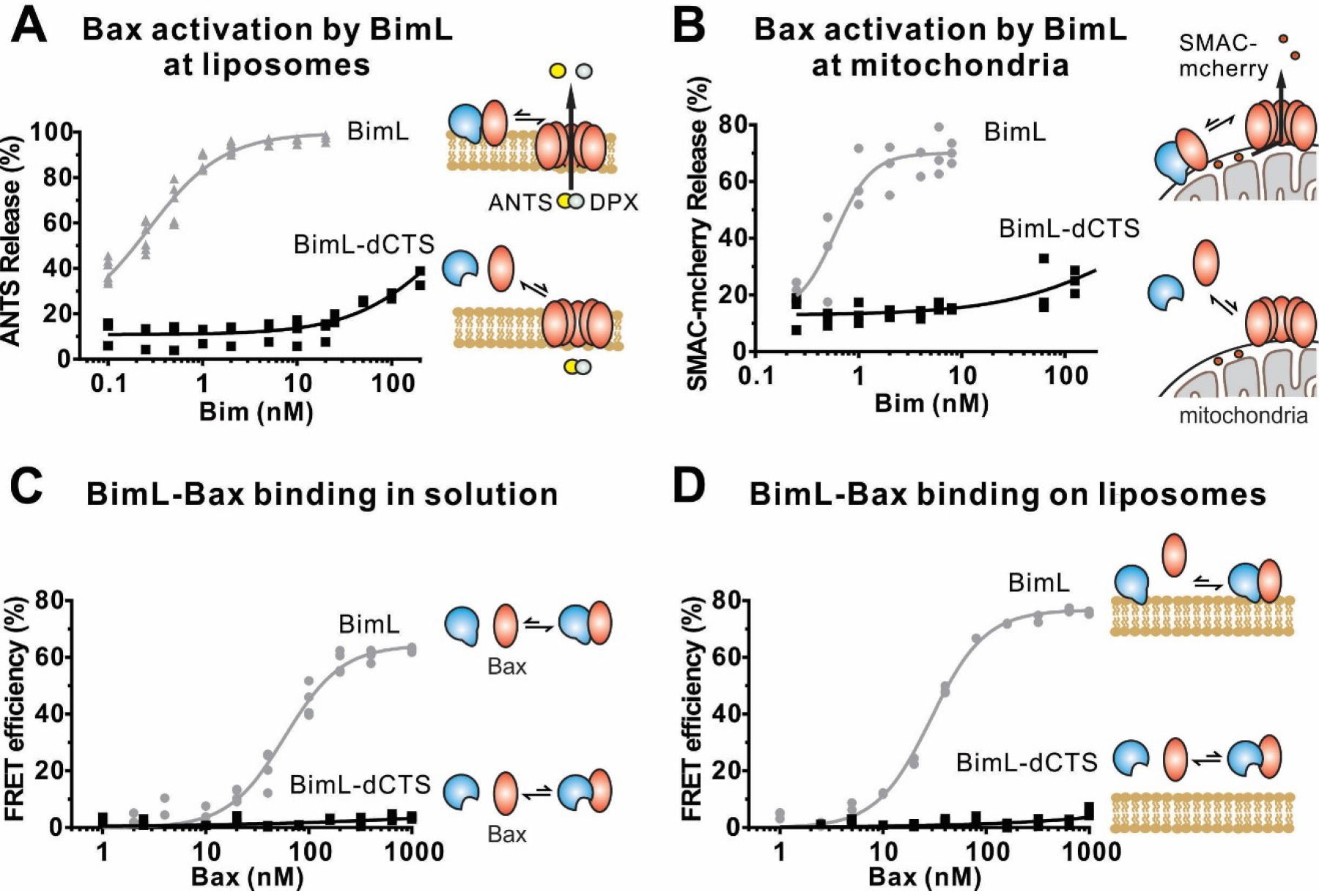
The Bim CTS is required to activate Bax to permeabilize membranes. Cartoons indicate the binding interactions being measured. Equilibria symbols indicate the predicted balance of complexes for BimL (blue), Bax (red) and membranes (tan). For each graph, data from three independent experiments are shown as individual points. Due to overlap, some points may not be visible. (A) Activation of Bax by BimL assessed by measuring permeabilization of ANTS/DPX filled liposomes (0.04 mg/mL) after incubation of Bax (100nM) with the indicated concentrations of BimL or BimL-dCTS. Fluorescence intensity, indicative of membrane permeabilization, was measured using the Tecan infinite M1000 microplate reader. (B) Permeabilization of the outer mitochondrial membrane by Bax (50 nM) in response to activation by the indicated amounts of Bim and BimL-dCTS was assessed by measuring SMAC-mCherry release from mitochondria. (C) Bim binding to Bax in solution measured by FRET. Alexa568-labeled BimL or BimL-dCTS (4 nM) was incubated with the indicated amounts of Alexa647-labeled Bax and FRET was measured from the decrease in Alexa568 fluorescence. (D) Bim binding to Bax measured by FRET in samples containing mitochondrial-like liposomes. FRET was measured as in (C) with 4nM Alexa568-labeled BimL or BimL-dCTS and the indicated amounts of Alexa647-labeled Bax.

To assess the impact of the Bim CTS on the interaction between Bim and Bax, binding was measured using Förster resonance energy transfer (FRET). For these experiments recombinant BimL proteins were labelled with the donor fluorophore Alexa568, while Bax was labelled with the acceptor fluorophore Alexa647. Unexpectedly, and unlike the BH3-only protein tBid (Lovell et al. 2008), BimL bound to Bax even in the absence of membranes (Figure 3C), while BimL-dCTS had no relevant Bax binding in the presence or absence of mitochondrial-like liposomes (Figure 3C-D). Binding of Bim to Bax in solution suggests that the CTS of Bim may be directly involved in Bim-Bax heterodimerization independent of Bim binding to membranes.

To confirm in our system that the labeled BimL proteins bind to membranes via the CTS sequence, binding of Alexa568-labeled recombinant BimL and BimL-dCTS to DiD labeled liposomes was measured by FRET (Figure 4A). In these experiments DiD serves as an acceptor for energy transfer from Alexa568 labeled BimL. The same approach was used to quantify BimL binding to mitochondrial outer membranes with mitochondria isolated from BAK^−/−^ mouse liver (Figure 4B), which lack Bax and Bak (Shamas-Din, Bindner, et al. 2013). In both cases, BimL spontaneously bound to membranes with picomolar affinity, while stable binding of BimL-dCTS to liposomes and mitochondria was not-detectable (Figure 4A-B). Furthermore, BimL-dCTS again had no relevant binding to Bax even in the presence of purified mitochondria (Figure 4C).

**Figure 4:**
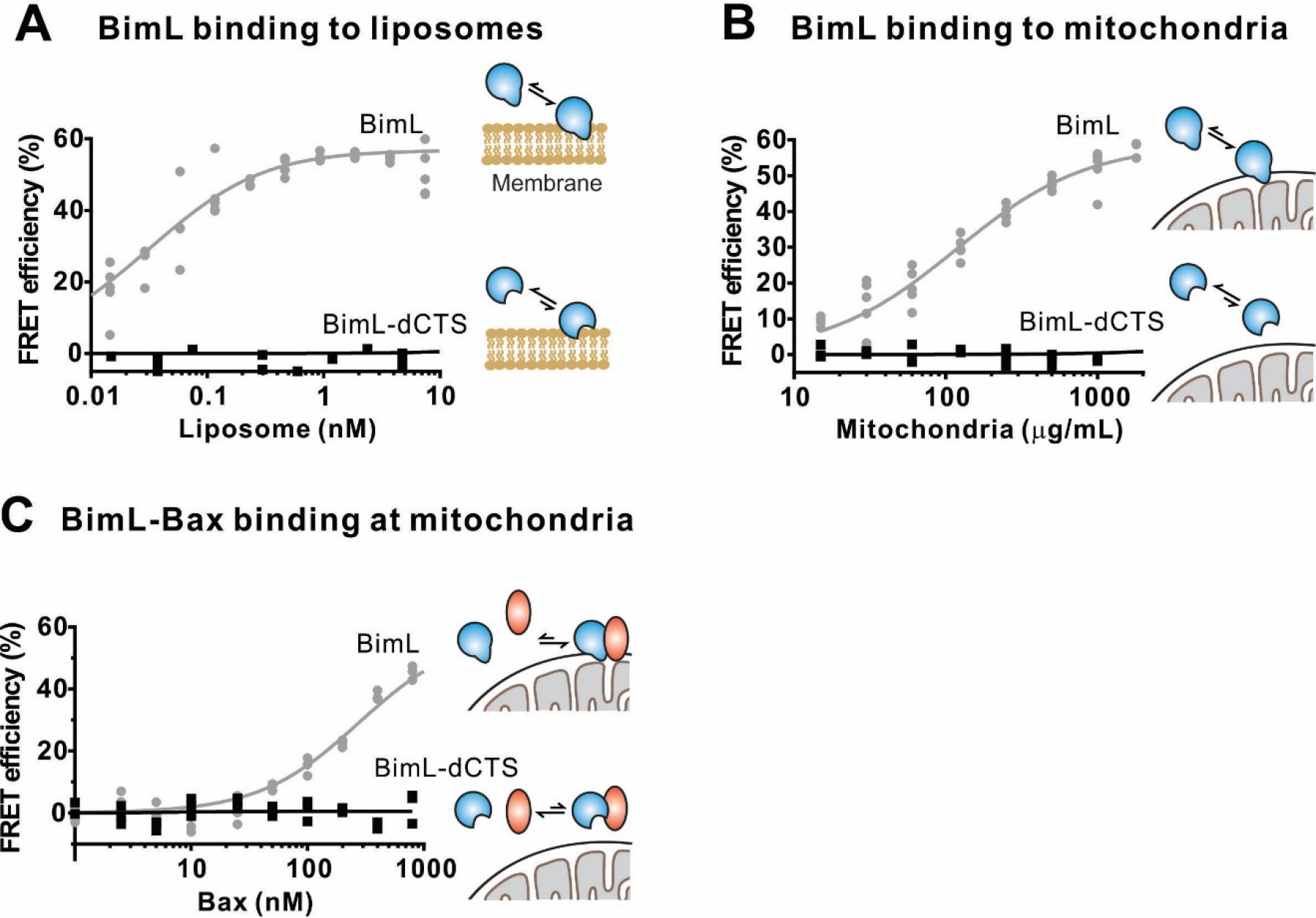
The Bim CTS is required to bind BimL to membranes *in vitro*. Cartoons indicate the binding interactions being measured. Equilibria symbols indicate the predicted balance of complexes for Bim (blue), Bax (red) and membranes (tan). For each graph, data from three independent experiments are shown as individual points. Due to overlap, some points may not be visible. (A) Bim binding to mitochondrial-like liposomes assessed by measuring FRET between 20nM Alexa568-labeled BimL or BimL-dCTS and the indicated amounts of DiD-labeled liposomes. (B) The CTS of Bim is necessary for Bim to bind to mitochondria. Binding of 4nM Alexa568-labeled BimL (n=5) or BimL-dCTS (n=3) to the indicated amounts of DiD labeled mouse liver mitochondria was assessed by measuring FRET. (C) Deletion of the CTS prevented Bim binding to Bax at mitochondria. Bim binding to Bax was measured by FRET in samples containing mouse liver mitochondria, 4nM Alexa568-labeled BimL (grey) or BimL-dCTS(black) and the indicated amounts of Alexa647-labeled Bax (n=3).

Taken together, our data strongly suggest that the CTS of Bim is required for both BimL to bind to membranes *in vitro* and for binding Bax with or without membranes. Alternatively purified BimL-dCTS may be completely non-functional. To demonstrate that purified BimL-dCTS binds to and inhibits Bcl-XL as shown for ^V^BimL-dCTS expressed in cells (Figure 1) and in (Liu et al. 2019), inhibition of Bcl-XL was measured using liposomes and mitochondria.

### The CTS is not required for BimL to inhibit Bcl-XL

In addition to direct Bax activation, Bim promotes apoptosis by binding to Bcl-XL and displacing either activator BH3-proteins (Mode 1) or activated Bax or Bak (Mode 2) (Llambi et al. 2011). In the ANTS/DPX liposome dye release assay, BimL-dCTS was functionally comparable to the well-established Bcl-XL inhibitory BH3-protein Bad in reversing Bcl-XL mediated inhibition of cBid (Figure 5A) or Bax (Figure 5B). Consistent with the observation that BimL-dCTS was less resistant to displacement by BH3 mimetics in live cells, in cell free assays BimL-dCTS was also less effective than BimL at displacing cBid or Bax from Bcl-XL (Liu et al. 2019). Nevertheless, when assayed with mitochondria BimL-dCTS disrupted the interaction between tBid and Bcl-XL resulting in Bax activation and permeabilization of mitochondria as measured by cytochrome c release (Figure 5C, solid black line). This activity is due to inhibition of Bcl-XL function, as in controls without Bcl-XL the same concentration of BimL-dCTS did not directly activate sufficient Bax to mediate MOMP (Figure 5C, dashed black line). Thus, purified BimL-dCTS is functional and can initiate MOMP by displacing direct-activators (Mode 1) or activated Bax (Mode 2) from Bcl-XL (Figures 5A-C). Finally, BimL-dCTS labelled with Alexa568 retained high affinity binding for Bcl-XL labelled with Alexa647 both in solution (Kd <16 nM) and on membranes (~35 nM apparent Kd on liposomes and on mitochondria) as measured by FRET (Figure 6B).

**Figure 5:**
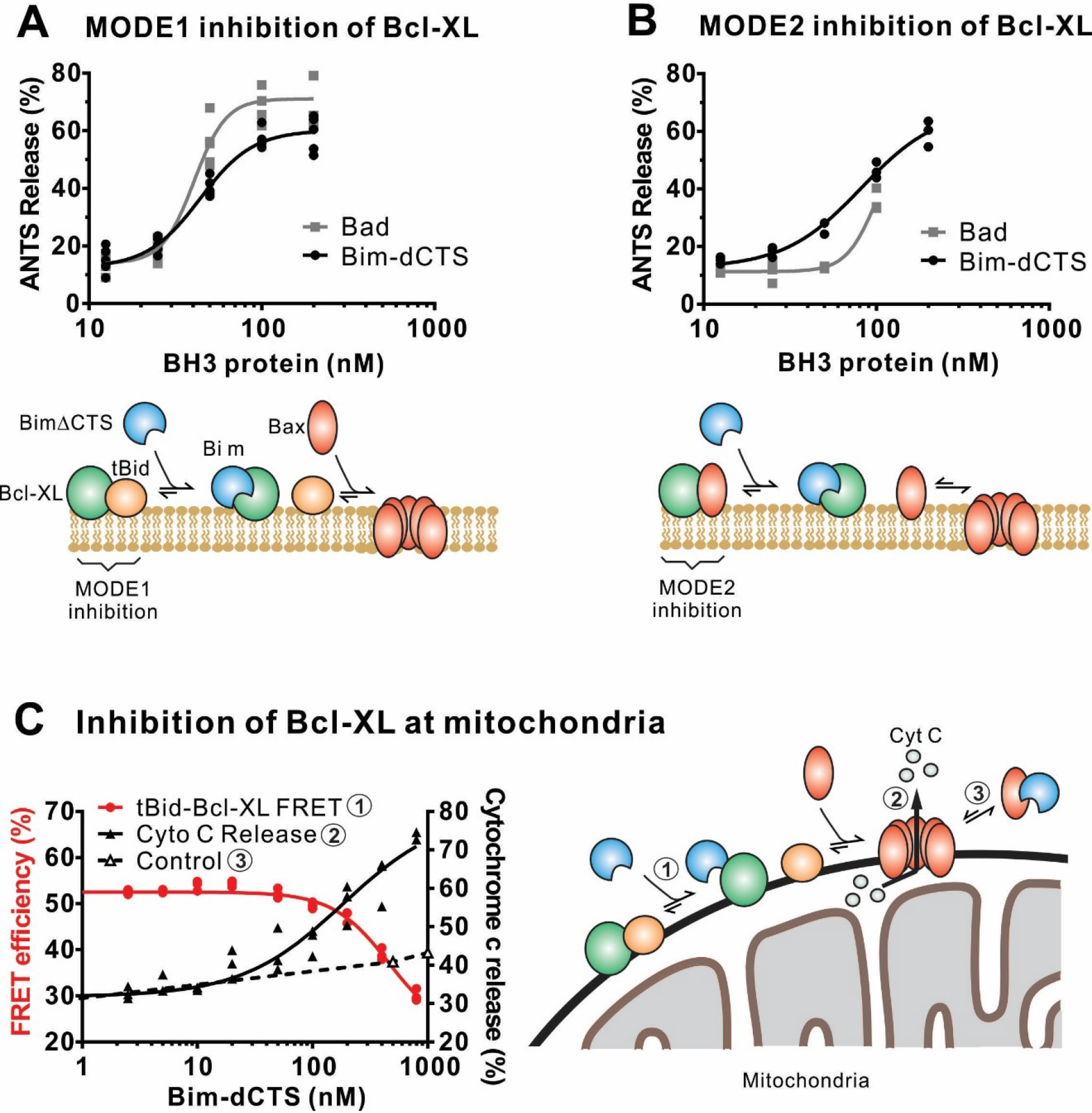
The Bim CTS is not required to inhibit Bcl-XL. (A-B) BimL-dCTS and Bad release tBid (A) or Bax (B) from Bcl-XL. 20nM tBid (A) or tBidmt1 (B), which activates Bax but does not bind Bcl-XL, were incubated with 100nM Bax, 40nM Bcl-XL, 0.04mg/mL ANTS/DPX liposomes, and the indicated amounts of either Bad or BimL-dCTS. Liposome permeabilization was assessed after incubation at 37°C for three hours by measuring the increase in fluorescence due to ANTS/DPX release. Cartoons indicate the interactions being measured, BimL (blue), Bax (red), tBid (orange), Bcl-XL (green), membranes (tan). (C) BimL-dCTS displaced tBid from Bcl-XL and permeabilized mitochondria. Mitochondria were incubated with Bcl-XL (40 nM), tBid (20 nM), Bax (100 nM) and mitochondria. Increasing concentrations of BimL-dCTS were added and displacement of tBid from Bcl-XL was measured by FRET. Mitochondria were then pelleted and cytochrome c release measured by western blotting. Control reactions containing only Bax and BimL-dCTS did not result in cytochrome c release (dotted line). Individual points are shown for three independent replicates. Not all points are visible due to overlap. The adjacent cartoon diagrams the interactions measured.

**Figure 6:**
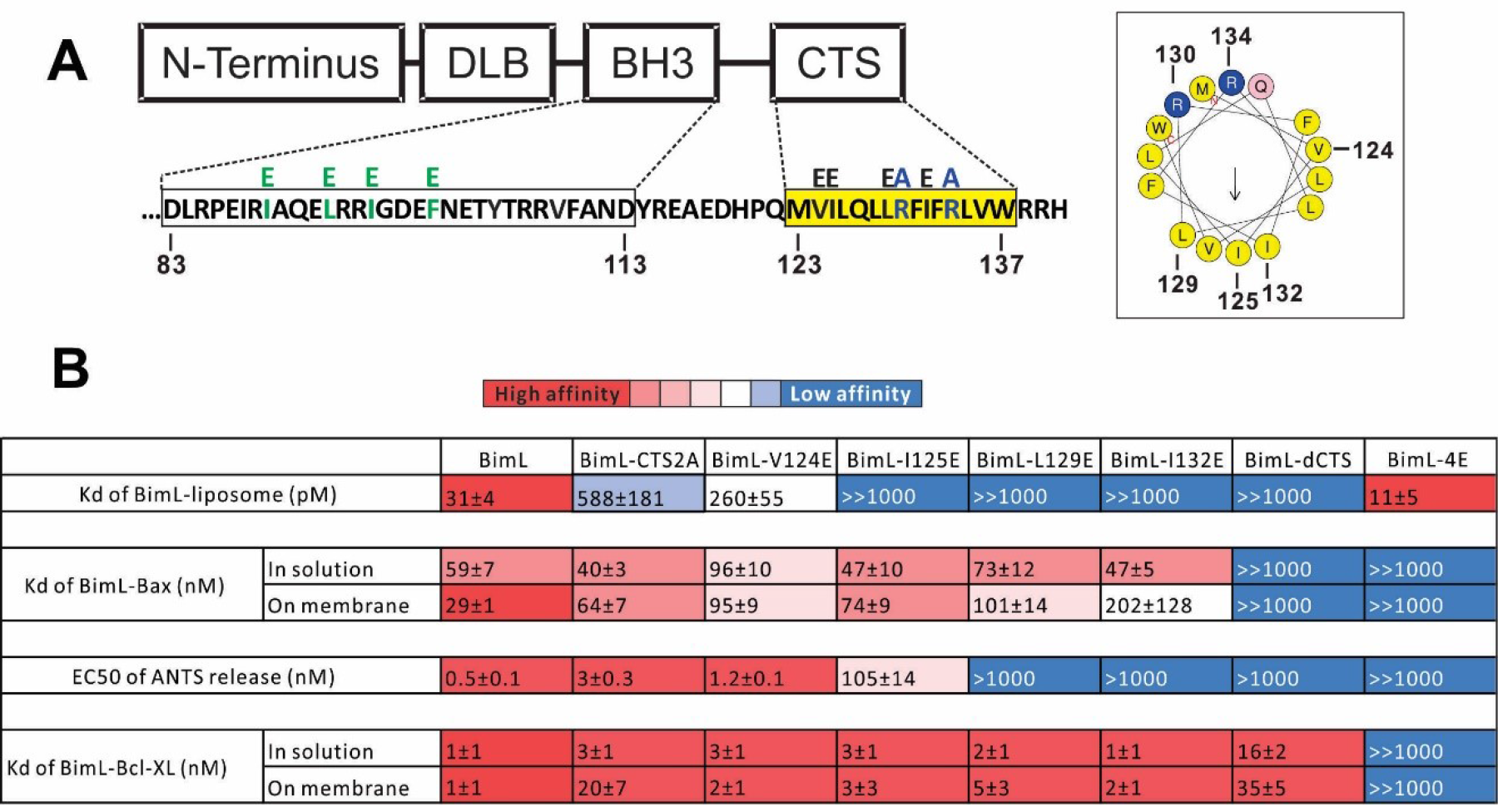
Residues within the Bim CTS distinctly regulate membrane binding and Bax activation. (A) Diagram of BimL depicting the various domains (DLB: dynein light chain binding motif) and the sequences of the BH3-domain and CTS. The four essential hydrophobic residues in BH3 domain that were mutated to glutamic acid are colored green. Two positive charged residues in the CTS mutated to alanine are colored blue. Glutamic acid mutations for individual hydrophobic residues in the CTS are indicated in black on top of the original sequence. A predicted alpha helix structure generated via HeliQuest software is shown on the right, indicating the amphipathic nature of the CTS. The arrow central to the helix shows the polarity direction for hydrophobicity. The Q indicated in pink is the last amino acid before the CTS. Other residues are colored as in the linear sequence. (B) Binding of BimL mutants to liposomes, Bax and Bcl-XL expressed as apparent dissociation constants (Kd) measured from raw data as in Figure 6 – figure supplement 1 for each binary interaction. Activation of Bax (EC50) measured from ANTS/DPX assays in Figure 6 – figure supplement 1. Values are mean ± SEM (n=3). The table is colour-coded in a heat map fashion as follows: red 0-40; light red 40-80; light pink 80-120; white 120-500; light blue 500-1000; Dark blue >1000. All values are nM except for binding to liposomes which is in pM. The Kds for ‘on membrane’ measurements are apparent values since diffusion for the protein fraction bound to membranes is in two dimensions while for the fraction of protein in solution diffusion is in three dimensions and several of the binary interactions take place in both locations. Apparent Kd values may also be affected by competing interactions with membranes.

### Different residues in the Bim CTS regulate membrane binding and Bax activation

To identify which residues in the Bim CTS mediate binding to membranes and/or Bax we generated a series of point mutations. Sequence analysis using HeliQuest software (Gautier et al. 2008) predicts that the Bim CTS forms an amphipathic α-helix (Figure 6A). Two arginine residues (R130&134) are predicted to be on the same hydrophilic side of the helix, whereas hydrophobic residues (e.g. I125, L129, I132) face the other side (Figure 6A). To determine the functional importance of these residues, Bim CTS mutants were created including: BimL-CTS2A in which R130 and R134 were mutated to alanine; and a series of single hydrophobic residue substitutions by glutamate (V124E, I125E, L129E, and I132E) (Figure 6A). To compare the effects of the CTS mutations on BimL binding interactions and function, we measured by FRET the Kds for the various binding interactions and the activities for the mutants to promote Bax mediated liposome permeabilization as EC50’s for ANTS release (Figure 6B and Figure 6 – Figure supplement 1A-E).

Mutation of individual hydrophobic residues on the hydrophobic side of the Bim CTS (BimL-I125E, BimL-L129E or BimL-I132E) abolished binding to membranes (Figure 6B). In contrast, mutations on the other side of the helix including BimL-V124E and BimL-CTS2A had less effect on Bim binding to membranes (Figure 6B). Despite the dramatic changes in affinity for membranes among Bim CTS mutants, the mutations did not abolish binding to Bax both in the presence and absence of membranes (Figure 6B). Indeed most of the mutants had Kd values for binding to Bax of less than 100 nM and to our surprise many of them bound to Bax better in solution than on membranes. This data further confirms that binding to membranes and Bax are independent functions of the Bim CTS. In the case of BimL-I125E, a mutant that activates Bax to permeabilize liposomes, the initial interaction with Bax must occur in solution as neither protein spontaneously binds to membranes (Figure 6B).

Unexpectedly, there was not a good correlation between BimL binding to membranes and Bax activation. For example, while BimL bound to membranes with a Kd of 31 pM, BimL-CTS2A and BimL-I125E bound to membranes very poorly (Kds of ~600 and >1000 pM, respectively) yet both mutants triggered Bax mediated membrane permeabilization, demonstrating that specific residues in the CTS rather than binding to membranes enabled BimL to mediate Bax activation. Moreover, BimL binding to Bax was also not sufficient to activate Bax efficiently. BimL-L129E and BimL-I132E are two Bim mutants that do not bind membranes, retain reasonable affinities for Bax in the presence of membranes (Kds ~100-200nM), but were unable to activate Bax (Figure 6B). These results indicate that these two residues play a key function in Bax activation. As expected, the negative control BimL-4E mutant does not bind to nor activate Bax even though its CTS is intact and the protein binds membranes (Figure 6B). This result confirms the essential role of the BH3 domain and suggests that the Bim CTS provides a secondary role in Bax binding rather than providing an independent high affinity binding site.

Both functional and binding assays for the various point mutants suggest that specific residues in the Bim CTS participate in Bim-Bax protein interactions that lead to Bax activation, however, these mutants did not clearly separate the membrane binding function of the CTS of Bim from a potential function in Bax activation. Thus, it remains possible that restoring membrane binding to BimL-dCTS would be sufficient to restore Bax activation function. To address this, we fused the mitochondrial tail-anchor from mono-amine oxidase (MAO residues 490-527, UniProt: P21397-1) to the C-terminus of BimL-dCTS to restore membrane binding with a sequence unlikely to contribute to Bax activation directly. This protein, BimL-dCTS-MAO, bound to purified mitochondria equally efficiently as full-length BimL (Figure 7A). However, while low nanomolar concentrations of BimL induced Bax-mediated SMAC-mCherry release from mitochondria, much higher concentrations of BimL-dCTS-MAO were required to trigger MOMP (Figure 7B). To analyze BimL-dCTS-MAO binding to Bax, Alexa568 labelled BimL-dCTS-MAO was incubated with Alexa647 labelled Bax in the presence (Figure 7C) or absence (Figure 7D) of purified mitochondria. The FRET results demonstrate that BimL-dCTS-MAO does not bind to Bax in solution (Figure 7D), however, restoring membrane binding to BimL-dCTS by adding the MAO sequence increased BimL-dCTS binding to Bax in the mitochondria marginally (Figure 7C, BimL-dCTS-MAO). Nevertheless, compared to BimL the binding of Bax by BimL-dCTS-MAO was at least an order of magnitude less efficient, explaining the reduced Bax activation function for BimL-dCTS-MAO (Figure 7B). Thus, similar to what was seen for Bim binding to Bcl-XL (Liu et al. 2019), specific sequences within the CTS of Bim increase Bim binding to Bax. However, consistent with sequences in the Bim CTS increasing the affinity of BimEL binding such that the heterodimer is resistant to disruption by BH3-mimetics (Liu et al. 2019), BimL-dCTS-MAO retained sufficient binding to Bcl-XL to functionally inhibit its sequestration of activated Bax (Figure 7E). By this definition BimL-dCTS-MAO functions as a sensitizer similar to the canonical sensitizer, Bad (Figure 7E).

**Figure 7:**
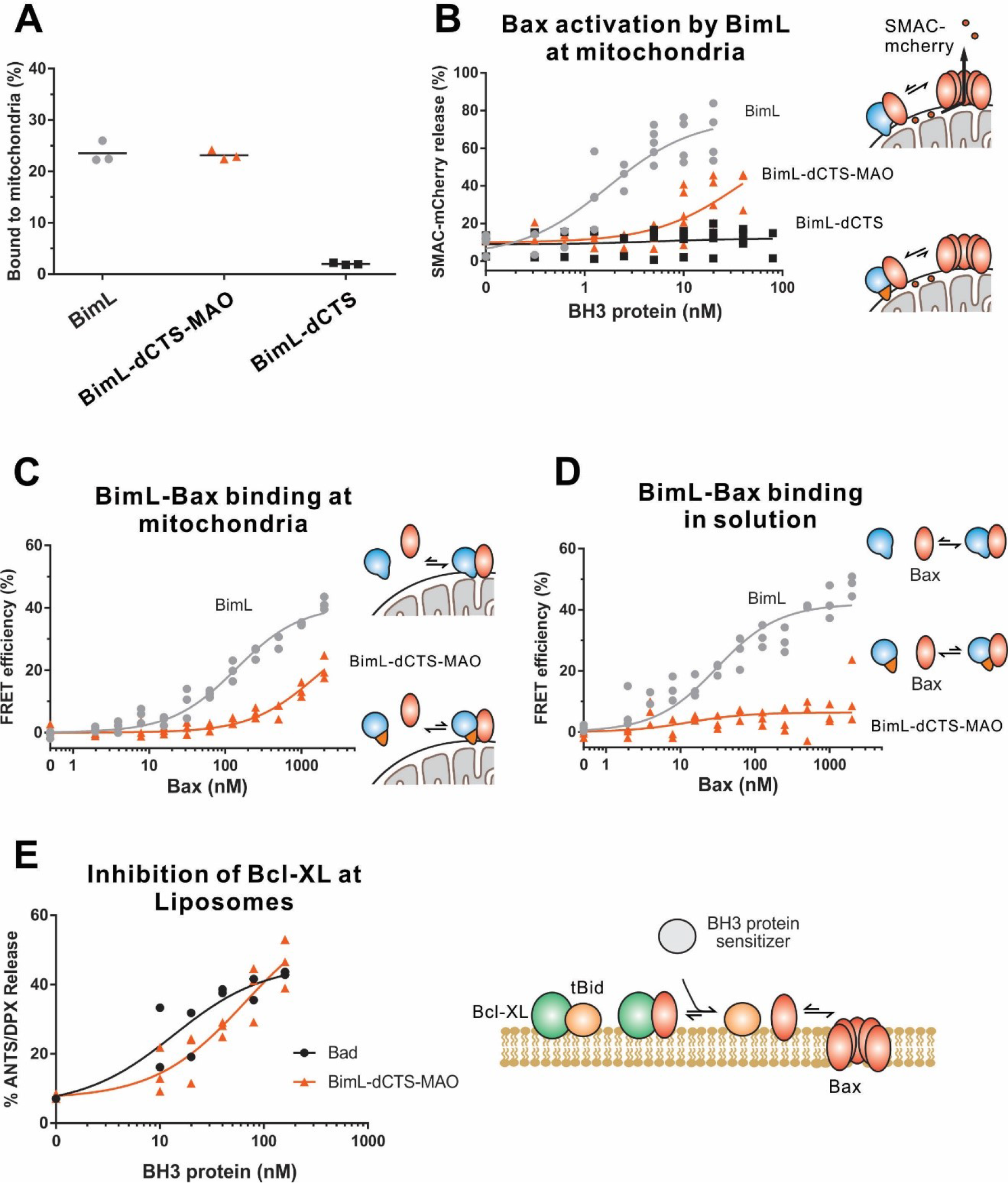
BimL-dCTS-MAO binds to mitochondria and Bax but activates Bax poorly. Cartoons at the side indicate the measurements being made with equilibria arrows representing the results obtained. Blue objects, Bim. Red ovals, Bax. Blue and orange objects, BimL-dCTS-MAO. (A) BimL-dCTS-MAO targets to mitochondria as efficiently as BimL. Alexa568 labelled single cysteine (Q41C) recombinant BimL, BimL-dCTS, and BimL-dCTS-MAO (20 nM) were incubated with 0.2 mg/ml of mitochondria purified from BMK Bax−/− Bak−/− cells for 40 minutes at 37°C. Mitochondria were pelleted by centrifugation for 10 minutes at 13000 x g. After correcting for background fluorescence binding to membranes was calculated from fluorescence signals in the supernatant and pellet fractions. (B) Restoring membrane binding to BimL-dCTS does not fully restore Bax activation function. Bax activation was measured by SMAC-mCherry release from purified mitochondria (n=5 for BimL and Bim∆CTS-MAO; n=3 for BimL-dCTS). The indicated amounts of BH3-only proteins were incubated with 20 nM of Bax in the presence of 0.2 mg/ml mitochondria. Reactions were incubated for 40 minutes at 37°C. (C-D) BimL-dCTS-MAO has reduced binding affinity to Bax compared to BimL in the presence (C) or absence (D) of mitochondria. 10 nM of Alexa 568-labeled BimL, BimL-dCTS or BimL-dCTS-MAO was incubated with the indicated amounts of Alexa 647-labelled Bax with or without of 0.2 mg/ml mitochondria. FRET was measured from the decrease in A568 fluorescence signal. (E) BimL-dCTS-MAO released activated Bax from sequestration by Bcl-XL.. ANTS/DPX filled liposomes were incubated with Bcl-XL (40 nM), cBid (20 nM), Bax (100 nM) to load Bcl-XL with activated Bax. Increasing concentrations of BimL-dCTS-MAO (orange line) or Bad (black line) were added and Bax-mediated liposome permeabilization was measured as an increase in ANTS fluorescence. Thus, BimL-dCTS-MAO functions as a sensitizer as illustrated in the adjacent cartoon.

### Residues within the Bim CTS are proximal to Bax in solution and on mitochondrial membranes

Our binding and mutagenesis data suggest that the Bim CTS binds to and activates Bax in solution and on membranes. To detect this binding interaction we used a photocrosslinking approach, in which a BimL protein was synthesized with a photoreactive probe attached to a single lysine residue positioned in the CTS using an *in vitro* translation system containing 5-azido-2-nitrobenzoyl-labled Lys-tRNA^Lys^ that incorporates the lysine analog (εANB-Lys) into the polypeptide when a lysine codon in the BimL mRNA is encountered by the ribosome. The BimL synthesized in vitro was also labeled by ^35^S via methionine residues enabling detection of BimL monomers and photoadducts by phosphor-imaging.

The radioactive, photoreactive BimL protein was incubated with a recombinant His6-tagged Bax protein in the presence of mitochondria isolated from BAK^−/−^ mouse liver lacking endogenous Bax and Bak to prevent competition and increase BimL-Bax protein interactions. Mitochondrial proteins were then separated from the soluble ones by centrifugation. Both soluble and mitochondrial fractions were photolyzed to activate the ANB probe generating a nitrene that can react with atoms in close proximity (≤ 12 Å from the Cα of the lysine residue). Thus, for photoadducts to form, the atoms of the bound Bax molecule are likely to be located in or near the binding site for the Bim CTS. The resulting photoadduct between the BimL and the His6-tagged Bax was enriched by Ni2+-chelating agarose resin and separated from the unreacted BimL and Bax monomers using SDS-PAGE. The ^35^S-labeled BimL in the photoadduct with His6-tagged Bax and BimL monomer bound to the Ni2+-beads specifically via the His6-tagged Bax or nonspecifically were detected by phosphor-imaging. A BimL-Bax specific photoadduct was detected when the ANB probe was located at four different positions in the Bim CTS on both hydrophobic and hydrophilic surfaces of the potential α-helix (Figure 8A). These photoadducts have the expected molecular weight for the Bim-Bax dimer, and were not detected or greatly reduced when the ANB probe, the light, or the His6-tagged Bax protein was omitted (Figure 8A). Consistent with the FRET-detected BimL-Bax interaction in both solution and membranes, the BimL-Bax photocrosslink occurred in both soluble and mitochondrial fractions. Less photocrosslinking occurred in the mitochondrial fraction likely due to the fact that in membranes homo-oligomerization of activated Bax competes with hetero-dimerization between BimL and Bax.

**Figure 8:**
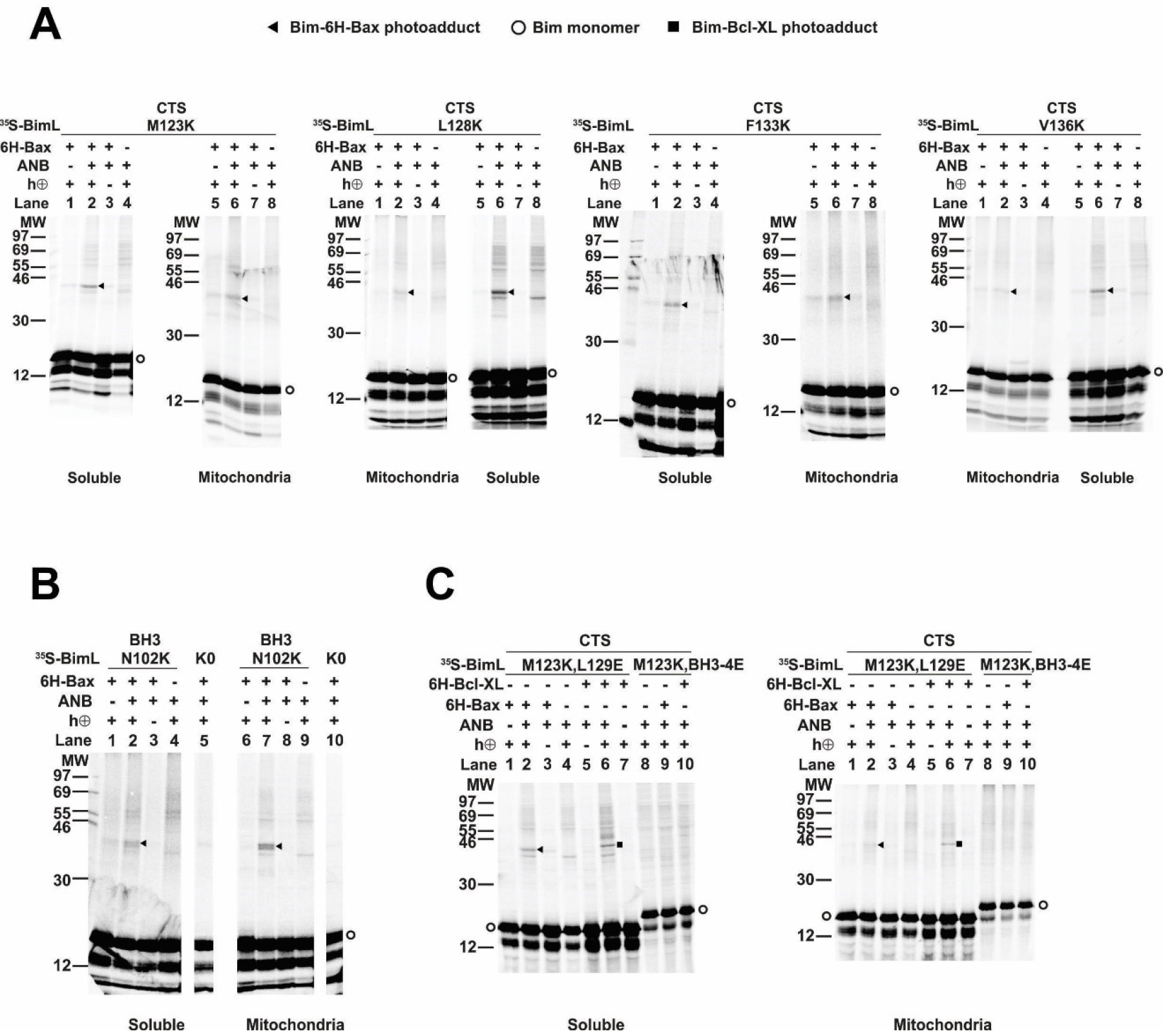
Residues within the Bim CTS interact with Bax. (A) Interaction of the Bim CTS with Bax in both soluble and mitochondrial fractions detected by photocrosslinking. The BimL proteins, each with a single εANB-lysine residue located at the position in the CTS indicated at the top of the panels and ^35^S-labeled methionine residues, were synthesized in vitro, and incubated with His6-tagged Bax protein (6H-Bax) in the presence of mitochondria lacking endogenous Bax since Bak. The mitochondria were then separated from the soluble proteins by centrifugation and both fractions were photolyzed. The resulting radioactive BimL/6H-Bax photoadducts were enriched with Ni^2+^-beads, and analyzed by SDS-PAGE and phosphor-imaging. BimL/6H-Bax dimer specific photoadducts were detected in both mitochondrial and soluble fractions and indicated by arrowheads. They were of reduced intensity or not detected in control incubations in which the ANB probe, light (hν) or 6H-Bax protein was omitted, as indicated. The radioactive BimL monomers are indicated by circles. The migration positions of protein standards are indicated by molecular weight (MW) in kDa. (B) Interaction of the Bim BH3 domain with Bax in both soluble and mitochondrial fractions detected by photocrosslinking. As a positive control illustrating the expected efficiency for the photocrosslinking experiments BimL protein with a single εANB-lysine located at the indicated position in the BH3 domain was used to photocrosslink 6H-Bax protein. In another control experiment, a lysine-null BimL protein that does not contain any εANB-Lys (K0) was used. As expected, a BimL/6H-Bax specific photoadduct was detected in the former but not the latter experiment. (C) The 4E mutation in the BH3 domain but not the L129E mutation in the CTS of Bim inhibited photocrosslinking of the Bim CTS to Bax. The BimL protein with either the BH3-4E or the L129E mutation and the εANB-Lys in the CTS was used in the photocrosslinking reaction with either 6H-Bax or 6H-Bcl-XL protein in both soluble and mitochondrial fractions. While the L129E mutation did not inhibit photocrosslinking of BimL to either 6H-protein, the BH3-4E mutation did. The BimL/6H-Bcl-XL photoadducts are indicated by squares.

As expected, BimL-Bax photocrosslinking was detected in both soluble and mitochondrial fractions when the ANB probe was positioned in the Bim BH3 domain as a positive control (Figure 8B). Crosslinking with the Bim BH3 domain is consistent with the canonical BH3 interaction well supported by experimental evidence including co-crystal structures and NMR models ((Walensky et al. 2008; Robin et al. 2015). Furthermore, loss of photocrosslinking for BimL mutants with the BH3 4E mutation that abolished binding to Bax demonstrates that direct binding between the proteins is required for crosslinking to be detectable (Figure 8C). Therefore, the crosslinking data suggests that similar to the BH3 domain, the Bim CTS binds to Bax. To further demonstrate that the CTS of Bim binds to Bax independent of both membrane binding and Bax activation the experiment was repeated with BimL-L129E, a mutant that binds Bax without activating it and that does not bind membranes (Figure 6B. As shown in Figure 8C, the L129E mutation in the CTS did not inhibit photocrosslinking of BimL to Bax in either the soluble or mitochondrial fractions. Furthermore, this mutant also resulted in photocrosslinking to Bcl-XL, consistent with data demonstrating that the Bim CTS also binds to this anti-apoptotic protein (Figure 8C and Liu et al. 2019).

### Bim CTS mutants that cannot activate Bax *in vitro* do not kill HEK293 cells

Together, our data suggests that specific residues within the Bim CTS are involved in different aspects of BimL functioning to activate Bax. Residue I125 is required for Bim to bind to mitochondria but is of lesser importance in activating Bax. In contrast, residues L129 and I132 are not required for BimL to bind Bax but are important for it to efficiently activate Bax. Finally BimL-dCTS functions only to bind and inhibit Bcl-XL. The defined mechanism(s) of these mutants makes them useful for probing the differential sensitivity of HEK293 and MEF cells to expression of ^V^BimL-dCTS as seen in Figure 1. Expression of the mutants in HEK293 cells by transient transfection revealed that similar to ^V^BimL-dCTS, expression of either ^V^BimL-L129E or ^V^BimL-I132E was not sufficient to kill HEK293 cells, despite expression of either mutant being sufficient to kill the primed MEF cell line (Figure 9). In contrast, HEK293 cells were killed by expression of ^V^BimL-I125E, albeit to a lesser extent than by ^V^BimL (Figure 9). This result is consistent with our findings with purified proteins showing that the EC50 for liposome permeabilization by BimL-I125E was 100 nM compared to ~ 1nM for BimL (Figure 6B). The activity of ^V^BimL-I125E also demonstrates that BimL binding to membranes is not required to kill HEK293 cells as BimL-I125E does not bind membranes (Figure 6B). Together, this data suggests that unlike MEF cells, only mutants of BimL that can efficiently activate Bax kill HEK293 cells.

**Figure 9:**
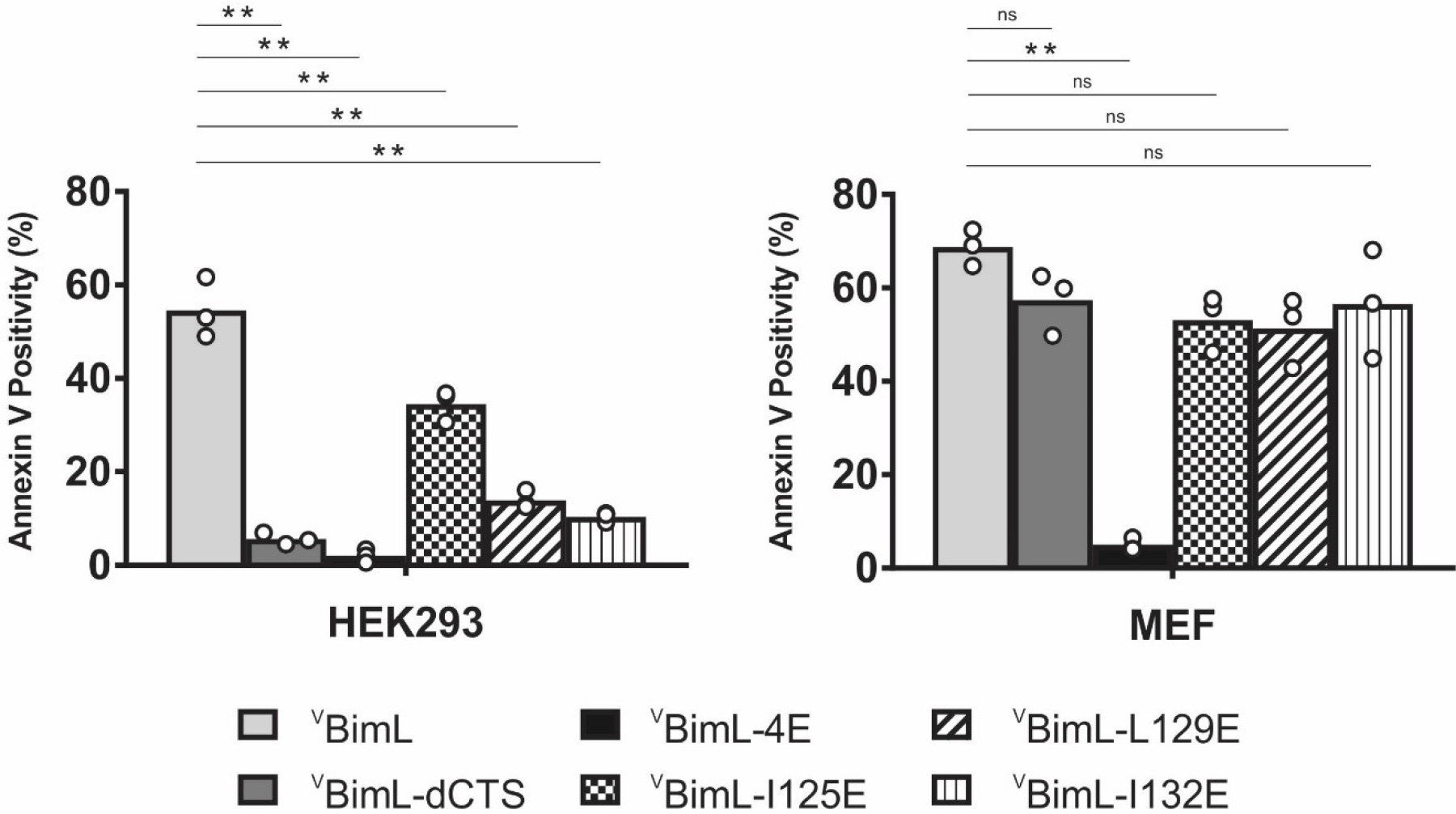
Bim CTS mutants that cannot activate Bax *in vitro* cannot kill HEK293 cells. The indicated cell lines were transiently transfected with DNA to express ^V^BimL, and the indicated ^V^BimL mutant proteins. Cells expressing Venus fusion proteins were stained with the nuclear dye Draq5 and rhodamine labelled Annexin V, and apoptosis was assessed by confocal microscopy as in Figure 1. The y-axis indicates Annexin V Positivity (%), which was calculated based on the total number of Venus expressing cells that also score positive for Annexin V rhodamine fluorescence. A minimum of 400 cells were imaged for each condition. Individual points (open circles) represent the average for each replicate, while the bar heights, relative to the y-axis, represent the average for all three replicates. A one-way ANOVA was used within each cell line followed by a Tukey’s multiple comparisons test to compare the means of each transfection group. *p-values less than 0.05, **p-values less than .01, ns, non-significant p-values (>0.05).

## Discussion

The apoptotic activity of Bim in live cells is likely mediated by a combination of functions that result in both activation of Bax and inhibition of anti-apoptotic proteins. Unlike any of the known BH3-proteins or small molecule inhibitors, BimL-dCTS inhibits all of the major multi-domain Bcl-2 family anti-apoptotic proteins without activating Bax or Bak. Thus expression of this protein in cells enables new insight into the importance of the extent to which a cell depends on the expression of anti-apoptotic proteins for survival (Figure 1A). Our results strongly suggest that the varying levels of apoptotic response of cell lines to BimL-dCTS reflect the extent to which that particular cell type is primed. Thus, HEK293 cells and adult neurons that are resistant to inhibition of Bcl-2, Bcl-XL and Mcl-1 but sensitive to activation of Bax, are functionally unprimed. Our data also reveal that intermediate states exist in which cells like CAMA-1 and BMK are partially resistant to inhibition of the multi-domain anti-apoptotic proteins (Figure 1A).

Partial resistance to expression of BimL-dCTS suggests that the flow of Bcl-2 family proteins between different binding partners leads to differential levels of dependency on the activation of Bax to trigger apoptosis. To illustrate this we have created a schema illustrating protein flow at the two extremes represented by fully unprimed and primed cells and the effects of mutations in the Bim CTS on regulating apoptosis (Figure 10). In the schema, flow is indicated by the different lengths of the equilibria arrows and illustrates the consequences of the various dissociation constants displayed in Figure 6B. BH3-proteins that do not efficiently activate BAX, such as BimL-L129E or BimL-I132E, interact primarily with anti-apoptotic proteins (illustrated here as Bcl-XL since it was possible to measure binding with purified proteins). The binding measurements in Figure 6B allow prediction of the outcome of more subtle differences in interactions for BimL and its mutants. For example, even though BimL-I125E activates Bax the concentration required is around 100nM while the dissociation constant for Bcl-XL is less than 3nM (Figure 6B) such that in cells BimL-I125E would preferentially bind and inhibit Bcl-XL rather than activate Bax (Figure 10B). While the CTS is necessary for Bim to activate Bax at physiologically relevant concentrations, membrane binding mediated by the CTS is not a prerequisite for interaction with Bax. Rather, binding to membranes increases subsequent Bax activation possibly through facilitating Bax conformational changes on the membrane (Figure 7B; compare BimL-dCTS-MAO and BimL-dCTS; and Figure 6B compare BimL, BimL-CTS2A and BimL-I125E). Thus it is likely that in cells expressing endogenous Bim, binding to membranes contributes to the efficiency with which the protein kills cells. Nevertheless, there exists the distinction in mechanism between Bim and tBid, as tBid requires membrane binding and a subsequent conformational change in order to bind and efficiently activate Bax (Lovell et al. 2008), while BimL can do so in solution via dual interactions by the Bim BH3 and CTS regions (Figure 3C and 10).

**Figure 10:**
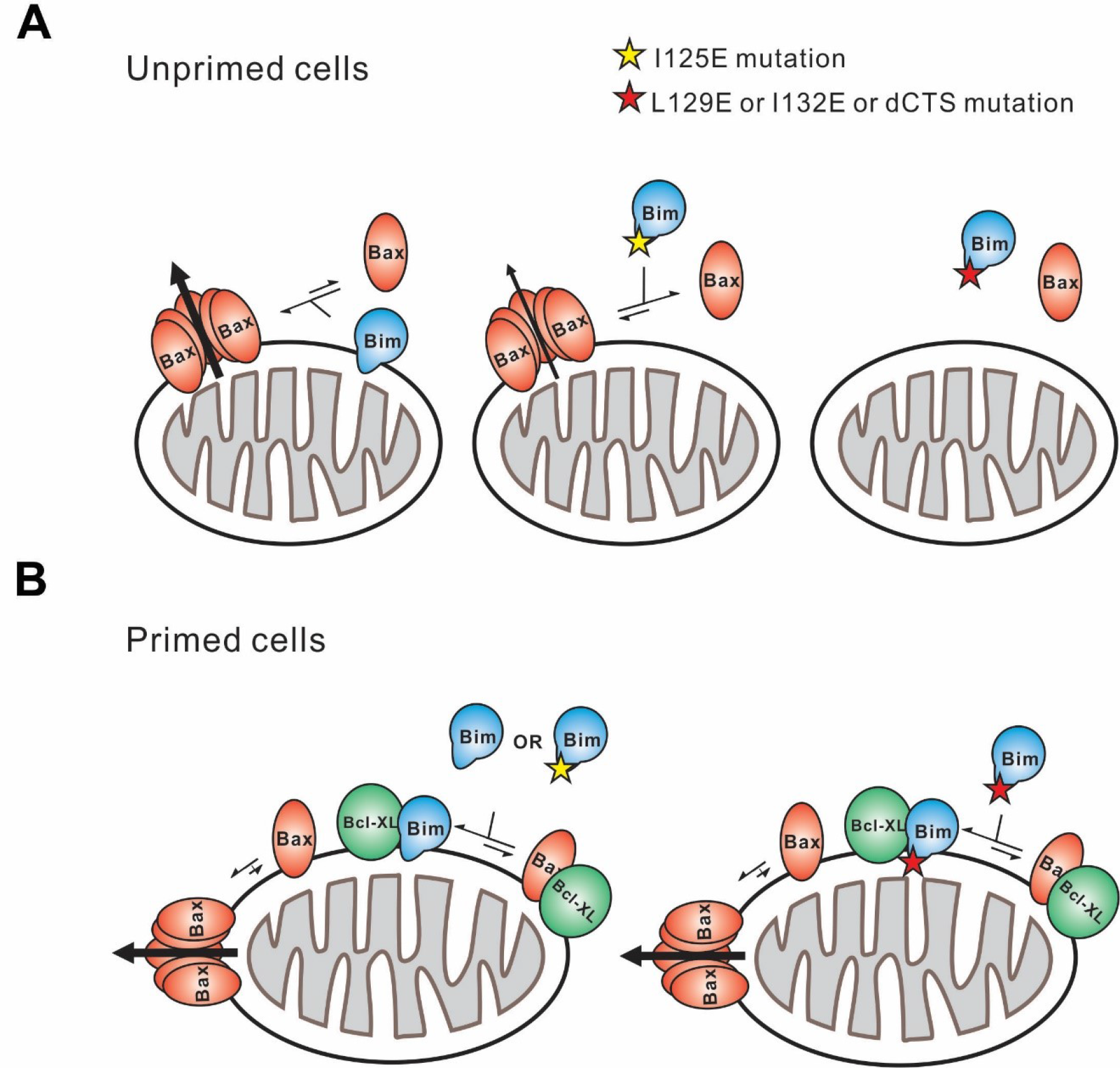
Schematic overview for the Bim CTS pro-apoptotic function. Interactions between BimL (blue), effector protein Bax (red) and Bcl-XL (green) are shown at mitochondria of unprimed and primed cells as indicated. Mutations of the Bim CTS are shown as a red or yellow star. Direction of protein flow into complexes indicated by lengths of the equilibria arrows is based on the Kds measured for the binding interactions (Figure 6B), the approximate cellular concentrations of the various proteins and activity assays with liposomes and mitochondria. (A) In unprimed cells the direct activation of Bax is the main function of Bim for inducing apoptosis. Comparison of BimI125E with BimL-L129E, BimL-I132E, and BimL-dCTS shows that the CTS, not membrane binding controls the activation of Bax by BimL (Figure 6B). BimL binds membranes and can activate Bax. BimL-I125E (yellow star) has no detectable membrane binding activity but still binds to and activates Bax, albeit with reduced activity compared to BimL (Figure 6B). At physiologically relevant concentrations BimL-L129E, BimL-I132E, and BimL-dCTS do not activate Bax. However, BimL-L129E and BimL-I132E binding to Bax is not reduced enough to account for the loss in Bax activation and membrane permeabilization suggesting these two residues are involved in activating Bax. (B) In primed cells, one or more pro-apoptotic proteins (activated Bax/Bak and/or a Bax-activating BH3-protein) are sequestered by anti-apoptotic proteins at the MOM. For simplicity only active Bax is shown. Depending on the amount of active pro-apoptotic protein sequestered and the amount of free inactive Bax and or Bak in the cell, BimL may initiate apoptosis primarily by inhibiting anti-apoptotic proteins or by activating Bax and inhibiting anti-apoptotic proteins. The Bim CTS is not required for binding to and inhibiting anti-apoptotic proteins as BimL-L129E, BimL-I132E, and BimL-dCTS bind to anti-apoptotic proteins such as Bcl-XL and release both pro-apoptotic BH3-proteins and Bax (Figure 5 and Figure 6B), thus enabling killing of primed cells.

The activities of the various Bim mutants analyzed here further suggest that specific residues in the Bim CTS enable physiological concentrations of Bim to activate Bax. That BimLV124E, BimLI125E and BimLL129E all bind Bax in solution and on membranes with similar affinities yet vary functionally to trigger Bax mediated liposome permeabilization by three orders of magnitude, suggests a specific role for this region in activation of Bax (Figure 6B) rather than the region simply increasing overall binding affinity. The situation is further complicated by another major role of the CTS of Bim in binding the protein to membranes. BimL-dCTS-MAO binds to mitochondria yet is defective in activating Bax to induce MOMP further suggesting a role for specific residues in the CTS binding to and activating Bax (Figure 7B). Such a role is consistent with our crosslinking data suggesting direct binding between these positions in the CTS of Bim and Bax (Figure 8) and that these residues particularly L129 (which corresponds to L185 in BimEL) increased the affinity for Bim binding to Bcl-XL such that it conferred resistance to BH3 mimetic drugs (Liu et al. 2019). Nevertheless, it remains formally possible that changes in binding affinity coupled with alterations in effective off-rate due to membrane binding may also contribute to the activation of Bax by Bim.

Currently, BH3-profiling is the technique used to assay the state of apoptotic priming for different tissue types, however, this technique requires the addition of BH3 peptides at high concentrations, and can only be performed on cells/tissues after permeabilization of the plasma membrane (Potter and Letai 2016). As an alternative, we propose lentiviral delivery and expression of BimL-dCTS be performed on living cells (or tissue samples), with readouts currently being used to assay cell death such as Annexin V staining, condensed nuclei, PI staining of nuclei, etc. Recently, it was reported in adults that most tissues are unprimed (Sarosiek et al. 2016), however the status of priming for different cell types that make up a single tissue may differ. In contrast, in tissue culture most cells are at least partially primed (Figure 1). We speculate that stress responses that result when fully or partially transformed cell lines are grown under non-physiological conditions (high glucose and oxygen, in the presence of serum and on plastic with abnormal stromal interactions) generally account for the dependence of these cell lines on continued expression of anti-apoptotic proteins. BH3-profiling can only provide an answer at the tissue level or for cell populations that can be isolated in sufficient quantities or easily cultured (Sarosiek et al. 2016). However, lentiviral delivery and expression of ^V^BimL-dCTS to specific cells in cultures, tissue slices and *in vivo* can provide the means to assay the level of dependence of individual cells on expression of anti-apoptotic Bcl-2 family proteins. This information could prove valuable to understanding which cell types may be most affected by small molecule BH3 mimetics used or in trial as chemotherapeutics and to better predict and prevent off-target toxicities that result in cell priming.

Overall, our data suggests a model in which the unusual CTS of Bim is not only required for binding to membranes but is directly involved in the activation of Bax. This function is crucial for BimL in killing unprimed cells. The CTS also increases the affinity of Bim for binding to Bcl-XL and Bcl-2 that is sufficient to induce apoptosis in primed cells (Figure 10). The very much higher affinity of Bim for Bcl-XL and Bcl-2 compared to Bax also ensures that in cells with excess anti-apoptotic proteins Bim is effectively sequestered and neutralized. In separate studies, we demonstrate that the additional affinity of the interaction of Bim with Bcl-XL and Bcl-2 provided by the Bim CTS is sufficient to dramatically reduce displacement of Bim by small molecule BH3 mimetics (Liu et al. 2019). Thus, regulation of apoptosis by Bcl-2 proteins is more complicated than presented in most current models. Moreover, the mutants and binding affinities described here provide the tools necessary for future studies of the relative importance of activation of Bax compared to inhibition of anti-apoptotic proteins in intact cells and in animals.

## Experimental Procedures

### Protein Purification

Wild type and single cysteine mutants of Bax, Bcl-XL, and cBid were purified as described previously (Kale et al. 2014). cBid mutant 1 (cBidmt1) was purified with the same protocol used for cBid (Kale et al. 2014). Bad was purified as described previously (Lovell et al. 2008).

His-tagged Noxa was expressed in E. coli strain BL21DE3 (Life Tech, Carlsbad, CA). E. coli cells were lysed by mechanical disruption with a French press. The cell lysate was diluted in lysis buffer (10mM HEPES (7.2), 500nM NaCl, 5mM MgCl_2_, 0.5% CHAPS, 1mM DTT, 5% glycerol, 20mM Imidazole) and Noxa was purified by affinity chromatography on a Nickel-NTA column (Qiagen, Valencia, CA). Noxa was eluted with a buffer containing 10mM HEPES (7.2), 300mM NaCl, 0.3% CHAPS, 20% glycerol, 100mM imidazole, dialyzed against 10mM HEPES 7.2, 300mM NaCl, 10% glycerol, flash-frozen and stored at −80 °C.

Purification of BimL and single cysteine mutants of BimL was carried out as previously described (Liu et al. 2019). Briefly, cDNA encoding full-length wild-type murine BimL was introduced into pBluescript II KS(+) vector (Stratagene, Santa Clara, CA). Sequences encoding a polyhistidine tag followed by a TEV protease recognition site (MHHHHHHGGSGGTGGSENLYFQGT) were added to create an in frame fusion to the N-terminus of BimL. All of the purified BimL proteins used here retained this tag at the amino-terminus. However, control experiments demonstrated equivalent activity of the proteins before and after cleavage with TEV protease. Mutations as specified in the text were introduced into this sequence using site-directed mutagenesis.

Bim was expressed in Arabinose Induced (AI) *E. coli* strain (Life Tech, Carlsbad, CA). *E. coli* were lysed by mechanical disruption with a French press. Proteins were purified from the cell lysate by affinity chromatography using a Nickel-NTA column (Qiagen, Valencia, CA). A solution containing 20mM HEPES pH7.2, 10mM NaCl, 0.3% CHAPS, 300mM imidazole, 20% Glycerol was used to elute the proteins. The eluate was adjusted to 150 mM NaCl and applied to a High Performance Phenyl Sepharose (HPPS) column. Bim was eluted with a no salt buffer and dialyzed against 10mM HEPES pH7.0, 20% glycerol, flash-frozen and stored at −80 °C.

### Protein labeling

Single cysteine mutants of Bax, Bcl-XL, cBid and Bad were labeled with the indicated maleimide-linked fluorescent dyes as described previously (Pogmore et al. 2016; Lovell et al. 2008; Kale et al. 2014). Single cysteine mutants of Bim were labeled with the same protocol as cBid with the exception that the labeling buffer also contained 4M urea.

### Bim binding to membranes

Liposomes (100 nm diameter) with a lipid composition resembling MOM were prepared as described previously (Kale et al. 2014). Mouse liver mitochondria were isolated from Bak^−/−^ mice as previously described (Pogmore et al. 2016). Liposomes and mitochondria were labeled with 0.5% and 2% mass ratios DiD, respectively (Life Tech, Carlsbad, CA). The single-cysteine mutant of Bim, BimQ41C, was labeled with Alexa568-maleimide and incubated with the indicated amount of unlabeled or DiD-labeled mitochondria or liposomes at 37° C for 1h. Intensities of Alexa568 fluorescence were measured in both samples as F_unlabeled_ and F_labeled_ respectively using the Tecan infinite M1000 microplate reader. FRET, indicating protein-membrane interaction, was observed by the decrease of Alexa568 fluorescence when Bim bound to DiD labeled membranes compared to unlabeled membranes. FRET efficiency was calculated as described previously (Shamas-Din, Bindner, et al. 2013). The data was fit to a binding model as described below. Lines of best fit were calculated using least squares in Graphpad Prism software.

### Membrane permeabilization

Membrane permeabilization assays with liposomes encapsulating ANTS and DPX were performed as described previously (Kale et al. 2014). To measure permeabilization of BMK mitochondria, the indicated amounts of proteins were incubated with mitochondria (1mg/mL) purified from BMK cells genetically deficient for Bax and Bak expressing mCherry fluorescent protein fused to the SMAC import peptide responsible for localization in the inter-membrane space. After incubation for 45 min at 37° C samples were centrifuged at 13000*g* for 10min to separate the pellet and supernatant fractions and membrane permeabilization was calculated based on the mCherry fluorescence in each fraction (Shamas-Din, Bindner, et al. 2013). For mouse liver mitochondria, cytochrome c release was measured by immunoblotting as described previously (Pogmore et al. 2016; Sarosiek et al. 2013).

### BH3 profiling

Heavy membranes enriched in mitochondria were isolated as described previously (Pogmore et al. 2016; Brahmbhatt et al. 2016). Membrane fractions (1mg/ml) were incubated with 500nM of the specified BH3 proteins (Bim, Bad and/or Noxa). For E15 brain mitochondria, 0.5mg/mL of membrane fractions were used and incubated with the indicated amounts of BH3-only proteins for 30 min at 37°C. Membranes were pelleted by centrifugation at 13000*g* for 10 min and cytochrome c release was analyzed by immunoblotting using a sheep anti-cytochrome c antibody (Capralogics). Mitochondria from embryonic mouse brains for BH3 profiling experiments were prepared from ~20 mouse embryos, E15 in age, following the same protocol used for liver mitochondria (Pogmore et al. 2016).

### Protein-protein binding

For FRET experiments, single cysteine mutants of cBid (126C), Bcl-XL (152C), Bax (126C), BimL (41C) and BimL mutants were purified and labeled with either Alexa 568-maleimide (donor) or Alexa 647-maleimide (acceptor) as specified. To determine binding constants donor protein was incubated with the indicated range of acceptor proteins and where specified liposomes or mitochondria. The intensity of Alexa568 fluorescence with unlabeled or Alexa647-labeled Bcl-XL was measured as F_unlabeled_ or F_labeled_ respectively and FRET was calculated as described in (Pogmore et al. 2016). All measurements were collected using the Tecan infinite M1000 microplate reader. Lines of best fit were calculated using least squares in Graphpad Prism software.

For each pair of proteins a dissociation constant (Kd) was measured in solution and with liposomes. Curves were fit to an advanced function taking into account the concentration of acceptor ([A]) change when [A] is close to Kd:

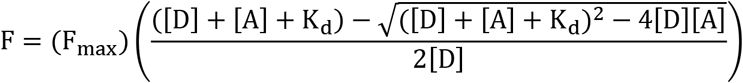

[D] is the concentration of donor, F indicates the FRET efficiency with the concentration of acceptor as [A], F_max_ is the maximum FRET efficiency in the curve (Pogmore et al. 2016).

### Photocrosslinking of Bim to Bax

The photocrosslinking method for studying interactions among the Bcl-2 family proteins has been described in detail (Lin, Johnson, and Zhang 2018). Briefly. [^35^S]Met-labeled BimL proteins with a single εANB-Lys incorporated at specific locations were synthesized using an in vitro translation system. 10 μl of the resulting BimL proteins were incubated at 37°C for 1 h with 1 μM of 6H-Bax or 6H-Bcl-XL protein and Bak^−/−^ mouse liver mitochondria (0.5 mg/ml total protein) in a 21-μl reaction adjusted by buffer A (110 mM KOAc, 1 mM Mg(OAc)_2_, 25 mM HEPES, pH 7.5). The mitochondrial and soluble fractions were separated by centrifugation at 13,000 x g and 4°C for 5 min, and the mitochondria were resuspended in 21 μl of buffer A. Both mitochondrial and soluble fractions were photolyzed to induce crosslinking via the ANB probe. The resulting samples were adjusted to 250 μl with buffer B (buffer A with 1% Triton X-100 and 10 mM imidazole) and incubated with 10 μl of Ni^2+^-chelating agarose at 4°C for overnight. After washing the Ni^2+^-beads three times with 350 μl of buffer B and one time with 400 μl of PBS, the photoadducts of the radioactive BimL protein and the 6H-tagged Bax or Bcl-XL protein and other proteins bound to the Ni^2+^-beads were eluted and analyzed by reducing SDS-PAGE and phosphor-imaging.

### Measurement of ell death in response to expression of ^V^BimL constructs

HEK293, CAMA-1, BMK, MEF, and HCT116 cells were maintained at 37°C (5% v/v CO_2_) in dMEM complete [dMEM, 10% Fetal Bovine Serum, 1% essential amino acids (Gibco, Grand Island, NY)]. Cells were seeded in CellCarrier-Ultra 384-well plates (1000 cells/well for BMK and MEF, 2000 cells/well for HEK293 and HCT116, 3000 cells/well for CAMA-1). One day later, cells were transfected using FugeneHD (Promega, Madison, WI) with plasmids encoding Venus, or Venus-fused BimL constructs in an EGFP-C3 backbone. Cell culture medium was added to each reaction (50μl/0.05μg DNA) and the whole mix added to each well (50μl/well) of a pre-aspirated 384-well plate of cells. After 24 hours, cells were stained with Draq5 and Rhodamine-labeled Annexin V and image acquisition was performed using the Opera QEHS confocal microscope (Perkin Elmer, Woodbridge, ON) with a 20x air objective. Untransfected cells and cells treated with 1μg/mL staurosporine were used as negative and positive controls for Annexin V staining. Cells were identified automatically using software as described previously (Shamas-Din, Bindner, et al. 2013). Intensity features were extracted using a script (dwalab.ca) written for Acapella high content imaging and analysis software (Perkin Elmer, Woodbridge, ON). Cells were scored as Venus or Annexin V positive if the Venus or Annexin V intensity was greater than the average intensity plus two standard deviations for the Venus or Annexin V channels in images of non-transfected cells. Cell death ascribed to the ^V^BimL fusion proteins was quantified as the percentage of Venus positive cells that were also Annexin V positive. For neuron cultures, cell segmentation using conventional methods could not be achieved due to complex cellular morphologies. Therefore, nuclei were first identified, then a ring region ~10% of nuclear area was drawn around each nuclei. Venus intensity was calculated for this ring region, representing the neuronal cell body, to determine if the neuron was expressing the Venus fluorescent protein.

Primary brain cortical neuron cultures were prepared from embryonic day 15, C57BL/6J mouse embryos as previously described (Mergenthaler et al. 2012). All animal breeding and handling was performed in accordance with local regulations and after approval by the Animal Care Committee at Sunnybrook Research Institute, Toronto. Briefly, after separation from hippocampus and subcortical structures, cortices were washed twice with ice-cold PBS, digested with 1x trypsin for 15 minutes at 37°C, washed twice with ice-cold PBS and then resuspended with a flame-treated glass pipette in N-Medium (DMEM, 10% v/v FBS, 2 mM L-glutamine, 10 mM Hepes, 45 μM glucose). The dissociated cortices were gently pelleted by centrifugation (200*g* for 5 minutes), N-media was removed, and neurons were resuspended and cultured in Neurobasal-Plus medium (ThermoFisher Scientific) supplemented with B27-Plus (ThermoFisher Scientific) and 1X Glutamax (ThermoFisher Scientific). Neurons were seeded at 5000 cells per well in a 384 well plate (Greiner μclear) after coating with poly-d-lysine (Cultrex). The medium was partially replaced on day five in culture with Neurobasal-Plus supplemented with B27-Plus and 1X Glutamax.

Lentivirus to express ^V^BimL and other BimL mutants were cloned into the pTet-O-Ngn2-Puro construct with the Ngn2 gene cut out. This construct was a kind gift from Dr. Philipp Mergenthaler, Charité Universitätsmedizin Berlin. Primary neuron cultures were infected with both ^V^BimL and rtTA lentiviral particles (~3μL of each concentrated stock) on the day of seeding. 24 hours later, Neurobasal-Plus medium containing lentiviral particles was removed and replaced with fresh Neurobasal-Plus medium.

Doxycyline (ThermoFisher Scientific) was added to 8 day *in vitro* old cultures of neurons at a concentration of 1μg/mL to induce ^V^BimL protein expression. 20 hours later, neurons were stained with 1 μg/mL Hoechst 33342 (Cell Signaling Technologies) and 1μg/mL propidium Iodide (Bioshop), then incubated for 30 min at 37°C. Confocal microscopy was performed immediately after.

### Lentiviral Production

Each lentivirus was made using the following protocol adhering to biosafety level 2 procedures. On day 0, lentiviral vectors psPax2 (10μg) and pMD2.G (1.25μg) were mixed with 10μg of desired ^V^BimL lentiviral construct in 1000μL of Opti-MEM media (ThermoFisher Scientific). Next, 42μL of polyethylenimine (PEI) solution [1mg/mL] was added, the mixture vortexed, then allowed to settle for 15 minutes at room temperature. After 15 minutes, 1.5×10^7^ of resuspended HEK293 cells and the transfection solution were mixed and seeded onto a 100mm culture dish with 10mL of dMEM complete plus 10μM of the caspase inhibitor Q-VD-Oph (Selleckchem), and left to incubate at 37°C (5% v/v CO_2_) for 72 hours. On day 3, media containing lentiviral particles was filter sterilized using a 0.45μm polyethersulfone filter, and mixed with polyethylene glycol (Bioshop) to achieve a final concentration of 10% (w/v). This was left to mix and precipitate the virus overnight at 4°C. On day 4, the media was centrifuged for 1 hour at 1600*g*, supernatant was then removed and the pellet was resuspended with 200μL of phosphate buffered saline. Resuspended virus was then stored at −80°C until needed.

## Acknowledgements

This work was funded by CIHR grant FRN 12517 to DWA and BL and CIHR Foundation grant FDN143312 to DWA, US NIH grants R01GM062964 and P20GM103640, OCAST grant HR16-026 and Presbyterian Health Foundation grant to JL. Q.L. held a post-doctoral fellowship from the Canadian Breast Cancer Foundation.

---

**Figure 1 – figure supplement 1.**
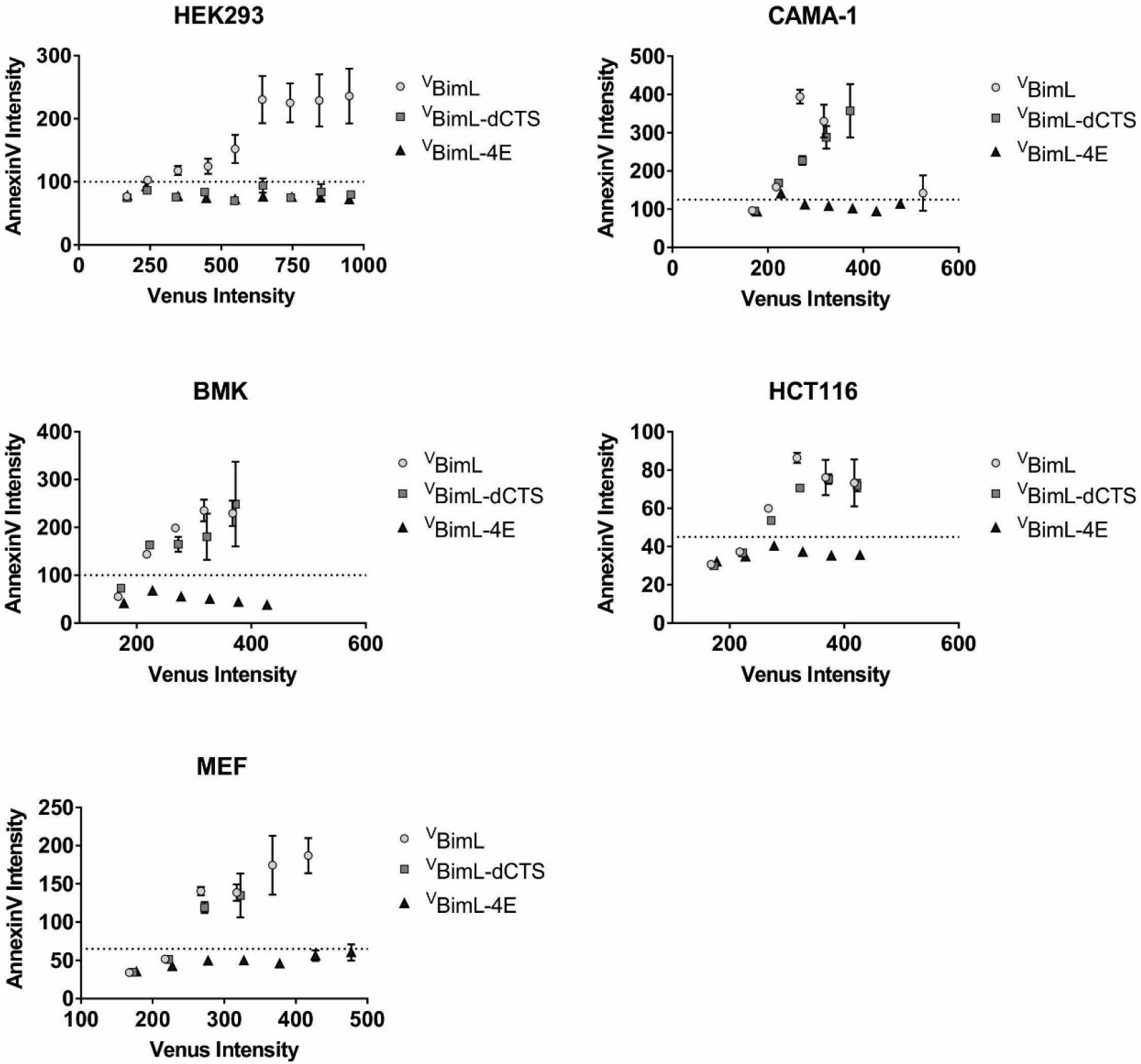
Correlation between expression of ^V^BH3-proteins and apoptosis measured as Annexin V labeling. For each cell line the Annexin V Rhodamine intensity was plotted against the Venus intensity. The Venus intensity acts as a surrogate for the relative amount of each of the BimL mutant proteins being expressed. Cells with intensity in the Venus channel equal to or less than the mean signal from untransfected cells were pooled as the first point, the subsequent points are data from intensity bins with 50 arbitrary unit increments. The horizontal dotted line indicates two standard deviations above the signal from untransfected cells in the AnnexinV channel. The cell line is indicated at the top of each panel.

**Figure S6 – Figure supplement 1.**
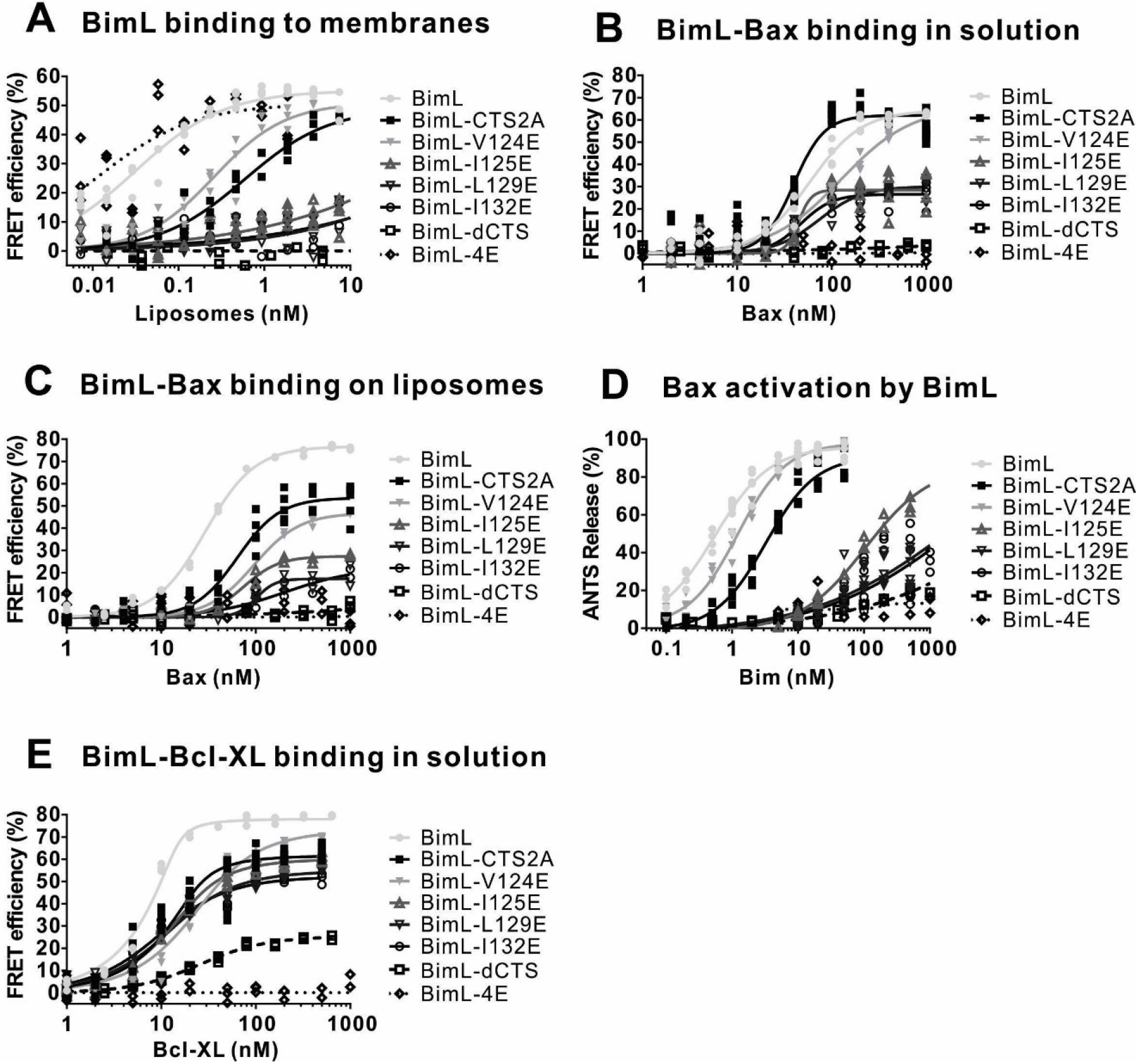
(A) BimL binding to membranes. (B) BimL binding to Bax in solution. (C) BimL binding to Bax with 2.9 nM liposomes. (D) Bax (100 nM) activation by BimL. (E) BimL binding to Bcl-XL in solution. For FRET experiments in (A-C) and (E), 20 nM of the indicated Alexa568-labeled BimL mutants (FRET donor) were incubated with the indicated concentrations of the Alexa647 labeled FRET acceptor labeled proteins. For each panel data from three independent experiments are shown as individual points, some points are not visible due to overlap. The mutants analyzed are indicated to the right of the graphs. To permit accurate estimation of the binding constants presented in Figure 6, data was collected to saturation for all mutants (for some curves 1600nM or 3200nM acceptor concentrations were required). For presentation purposes all curves were truncated at 1000 nM.

